# Rational engineering enhances the signal and modularity of an RNA barcoding technology to track gene transfer in microbiomes

**DOI:** 10.64898/2026.04.20.719664

**Authors:** Lavanya Karinje, Jonathan J. Silberg, James Chappell, Lauren B. Stadler

## Abstract

Horizontal gene transfer drives microbial evolution and offers a powerful strategy for precision microbiome engineering. To track gene transfer within complex communities, we previously developed RNA-Addressable Modification (RAM), an RNA-barcoding technology where a mobile catalytic RNA barcodes host 16S ribosomal RNA (rRNA) upon gene transfer. However, the first-generation RAM suffered from low barcoding efficiency and lacked modularity, limiting its sensitivity and versatility. Here, we present RAM v2, a re-engineered system with significantly enhanced performance and modularity. By incorporating natural ribozyme structural motifs and improved barcode stability, we achieved a ∼200-fold increase in barcoded rRNA signal. To enhance modularity, we integrated CRISPRi-based repression and ribozyme insulators, facilitating easy promoter swapping. We validated RAM v2 on a mobilisable plasmid delivered to a complex wastewater microbial community, demonstrating a substantial increase in signal over the original system while barcoding similar taxa. These improvements enable higher-resolution, more sensitive monitoring of horizontal gene transfer, providing a robust toolkit for accelerating the study of gene transfer in microbial communities and advancing targeted microbiome engineering.

## INTRODUCTION

Horizontal gene transfer (HGT) is a primary driver of microbial evolution, occurring through mechanisms such as transformation, phage transduction, and conjugation (1, 2). These processes allow microbes to rapidly acquire new functions, including antibiotic resistance (3), metabolic functions (4), and phage resistance (5). By facilitating the acquisition of complex and novel traits, HGT enables microbes to adapt to environmental stressors on a timescale far shorter than that required for the accumulation of beneficial mutations (6). HGT is also critical for delivering DNA into individual microbes and microbial communities (7–9). It can be harnessed to spread genes in a microbiome for therapeutic applications, such as degradation pathway genes for bioremediation (10), and biosensors to record environmental exposure to harmful chemicals (11).

Despite its importance, measuring HGT *in situ* within microbial communities remains a significant challenge. While genetically encoded reporters, such as fluorescent proteins or antimicrobial resistance genes, have been used to track gene transfer (12–14), they have several inherent limitations. These reporters rely on correct translation and protein folding, may depend on culture-based selection, and can impose a significant metabolic burden on the host cell (15, 16). As a result, these methods may not accurately reflect the full extent of gene transfer and may yield false positives and negatives due to autofluorescence, native antibiotic resistance, and variation in reporter expression. Preparation of samples to detect these types of reporter outputs can also be laborious, limiting throughput. In addition, genomic diversification can be driven by the rare biosphere (17), yet monitoring gene transfer within these low-abundance populations is especially challenging.

To address these challenges, we developed a novel reporter of gene transfer called RNA-Addressable Modification (RAM). RAM uses an engineered ribozyme with a designable guide RNA to bind and splice an RNA barcode onto host 16S ribosomal RNA (rRNA) at a conserved uracil (U) within a highly conserved rRNA motif (18) (**Figure 1A**). By embedding RAM into mobile DNA, when this DNA enters a cell, the catalytic RNA (cat-RNA) adds a barcode onto 16S rRNA to record gene transfer. Barcoded rRNA can then be isolated, amplified, sequenced, and analysed to identify who participated in gene transfer. For recording gene transfer in microbial communities, RAM offers several potential benefits. As the process of recording gene transfer occurs at the transcriptional level, it eliminates the need for translation and thus offers a potentially broader host range (19). By functioning independently of the host translational machinery, RAM also minimises the fitness cost typically associated with protein-based reporters (15, 16). RAM also enables culture-independent monitoring, a critical capability for analysing the large number of unculturable strains in microbial communities. As RAM is read out using established, targeted sequencing techniques (e.g., 16S rRNA amplicon sequencing), it is readily accessible, leverages existing analysis pipelines, and offers a high degree of sensitivity. Finally, because RAM appends a barcode to report on gene transfer, barcodes corresponding to different DNA types or mobile genetic elements can be multiplexed to simultaneously map their host ranges or study their interactions in a single experiment. To date, RAM has been used to study plasmid conjugation from *Escherichia coli* and transduction by bacteriophage P1 in wastewater communities, revealing substantially broader host ranges for these plasmids and bacteriophage than previously recognised (18, 20).

**Figure 1.**
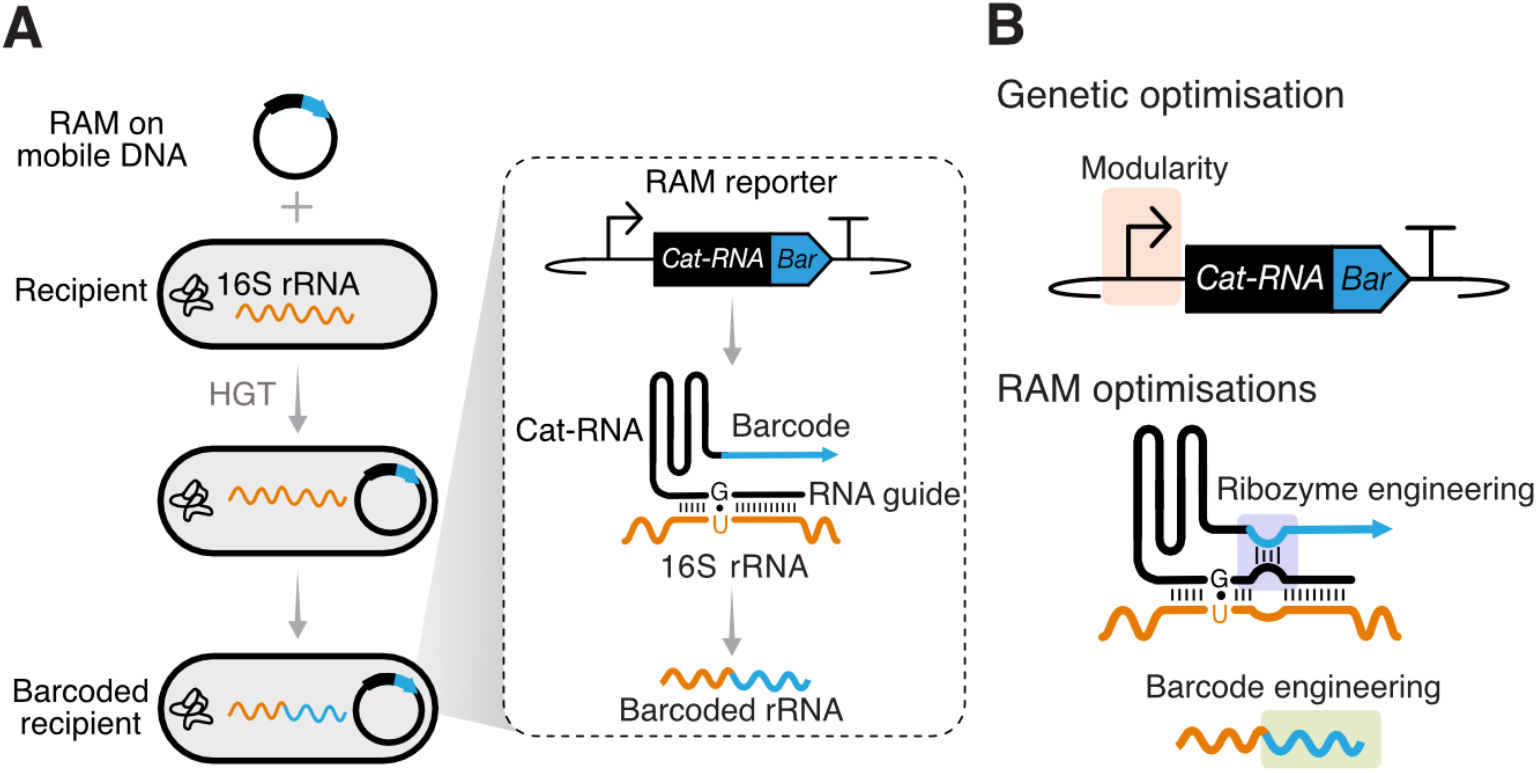
Mechanism and rational engineering of RNA Addressable Modification (RAM), a genetic reporter of horizontal gene transfer. (**A**) RAM uses an engineered catalytic RNA that splices a synthetic RNA barcode onto host 16S rRNA. By embedding the RAM reporter onto mobile DNA, it can be used to measure uptake and identify participants in horizontal gene transfer (HGT). (**B**) Overview of the rational optimisations to enhance the modularity of the RAM genetic cassette and the RAM signal by ribozyme and barcode engineering.

While RAM provides an innovative and accessible technology for measuring gene transfer, the first-generation system has several limitations and considerable room for improvement. In particular, the cat-RNA and barcode sequences were not optimised to maximise splicing efficiency or barcoded rRNA signal. Consequently, cat-RNA splicing efficiency was relatively low; for example, in *E. coli*, only ∼0.01% of 16S rRNA is barcoded under exponential growth conditions. This low barcoding efficiency limits the overall RAM signal and reduces sensitivity for detecting HGT events. Additionally, the original genetic construct offered limited modularity, preventing the facile exchange of genetic parts such as promoters. This is due to the dependence on a specific promoter and repressor pair (Pcym and CymR), which are used to repress the transcription of RAM in donor cells during conjugation measurements. This limits flexibility to modify promoter strength, expand the host range by using broad-host-range promoters (19), or enhance RAM transcription in specific taxonomic groups (e.g., Gram-positive bacteria) (21).

To address these limitations, we rationally engineered the RAM reporter and genetic construct to enhance the signal output and modularity (**Figure 1B**). First, we engineered the ribozyme to mimic tertiary interactions present in the native ribozyme (22), thereby improving splicing efficiency by ∼4-fold. We then rationally engineered the barcode of RAM to enhance the stability of the spliced product by mimicking the structure of natural tRNAs (23) and optimising the sequence to reduce RNase-mediated decay (24, 25). When combined with the optimised ribozyme, these new versions of RAM, termed RAM version 2 (RAM v2), increased the RAM signal ∼200 fold compared to the original RAM. We then established a modular RAM genetic design that leveraged CRISPR interference (CRISPRi) repression (26) and genetic insulating ribozymes (27), enabling facile and predictable promoter exchange. Finally, we validated the use of RAM v2 for measuring HGT in communities by measuring conjugation of a plasmid from an *E. coli* donor into a complex wastewater community.

## RESULTS

### Rational engineering of ribozyme interactions improves splicing efficiency and RAM signal

To improve RAM signal output, we first focused on rationally engineering the catalytic RNA to improve its splicing efficiency. The wild-type *Tetrahymena thermophila* group I intron splicing ribozyme has various tertiary interactions to facilitate splicing of the flanking exons (22, 28). One of these interactions occurs between the loop of the paired region 1 (P1) and the 3’ exon to transiently form a paired interaction called P10, which aligns the 3′ exon for the second transesterification step of catalysis (Supplementary Fig. 1). Prior work on engineering trans-splicing ribozymes has shown that replicating this interaction by including a bulge within the guide-target RNA duplex can enhance splicing efficiency (22). However, this was absent in our original RAM design (RAM v1). To test if including this would enhance RAM signal, we reintroduced this interaction in two different designs: (i) by adding the sequences present in the wild-type ribozyme into the P1 bulge and barcode (i.e., the 3’ exon) to reform the native P10 (WT P10); and (ii) by adding the P1 bulge to complement four nucleotides of the barcode to form a synthetic P10 (Syn. P1; Fig. 2A). To test these designs, the modified cat-RNAs with the P10 interactions were separately expressed from plasmids in *E. coli* using a constitutive promoter. Total RNA was extracted, and the barcoded 16S rRNA and native 16S rRNA were quantified by RT-qPCR (Fig. 2B and Supplementary Fig. 2). These experiments revealed that both RAM versions that included the P10 produced 3-5 fold higher levels of barcoded 16S rRNA than RAM v1. As there was no significant difference in barcoded 16S rRNA between WT P10 and Syn. P10 designs, we chose the Syn. P10 design for subsequent experiments. Taken together, these results revealed that the RAM splicing efficiency could be improved by rationally incorporating transient RNA interactions present in the native ribozyme.

**Figure 2.**
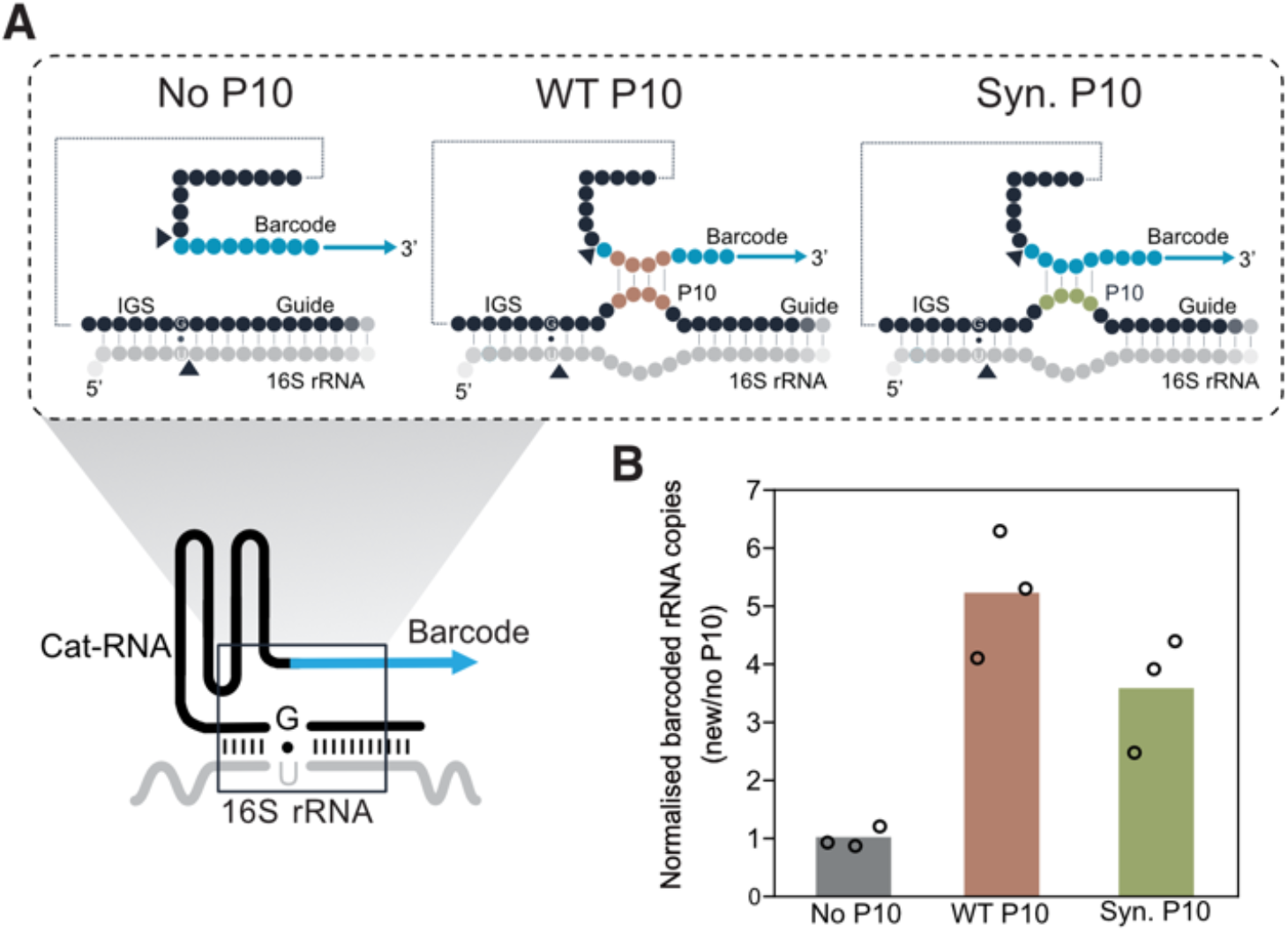
Rational engineering of the cat-RNA P10 interaction increases barcoding efficiency in *E. coli*. (**A**) Schematic of the cat-RNA, which targets conserved sequences within 16S rRNA for trans splicing reactions. Three different designs are shown, including the original RAM version 1 (v1) design (no P10) and two designs rationally engineered to allow for a transient interaction called P10 between the barcode and the ribozyme at the P1 bulge. (Middle panel) The cat-RNA is engineered to contain the native P10 interaction from the wild-type *T. thermophila* ribozyme (WT P10), and (right panel) the cat-RNA is engineered to contain a P1 bulge complementary to the existing barcode (Syn. P10). (**B**) Quantification of barcoded 16S rRNA using RT–qPCR in *E. coli* cells containing plasmids encoding RAM v1 (no P10) or the new modified versions. Barcoded 16S rRNA was normalised to native 16S rRNA. RT–qPCR data was normalised to the ‘No P10’ design. Individual data points show three biological replicates, and bars show the mean. The WT P10 and Syn. P10 signals are significantly higher than the No P10 version (P<0.022 for both; *t-*test).

### Barcode structure and length impact stability and signal

Beyond splicing efficiency, we reasoned that the RAM signal is also set by the stability of the barcoded 16S rRNA product, whose degradation rate determines its steady-state abundance per cell. We hypothesised that the stability of the RAM product could be improved by altering the sequence and structure of the barcode. In the original RAM design (RAM v1), a non-coding 518-nucleotide fragment of GFP was used as a barcode. To investigate if this barcode could be optimised, we explored two distinct designs. The first design, called the tRNA barcode, uses an engineered tRNA scaffold that has been widely used as an expression platform to maximise RNA expression in cells and in vitro by preventing RNase degradation (23). Specifically, we used *E. coli* methionine tRNA and replaced the anti-codon loop with a 28-nucleotide RNA sequence. This sequence was included to serve as a site for reverse transcription primer binding and to allow for the inclusion of indexes for creating multiplexable barcodes. The second design, called the minimised barcode, aimed to minimise barcode length while maximising secondary structures and avoiding well-known RNase E cleavage sites, such as GUAUUU (29, 30). This design was guided by recent high-throughput screens of synthetic mRNA stability, which identified these as major drivers of RNA half-life (24, 31) (Fig. 3A). These two barcode designs were cloned onto the Syn. P10 version of the cat-RNA, the plasmids were transformed into *E. coli* cells, and the native and barcoded rRNA measured by RT-qPCR (Fig. 3B, Supplementary Fig. 3 and 4). Compared to the RAM v1, this new design resulted in a ∼200-fold increase in RAM signal. We also confirmed that these barcodes increased RAM signal independently of the ribozyme containing P10 interactions and observed a ∼75-fold increase relative to the original barcode (Supplementary Fig. 5). These results show that the RAM signal can be dramatically improved through rational engineering of the RAM barcode.

**Figure 3:**
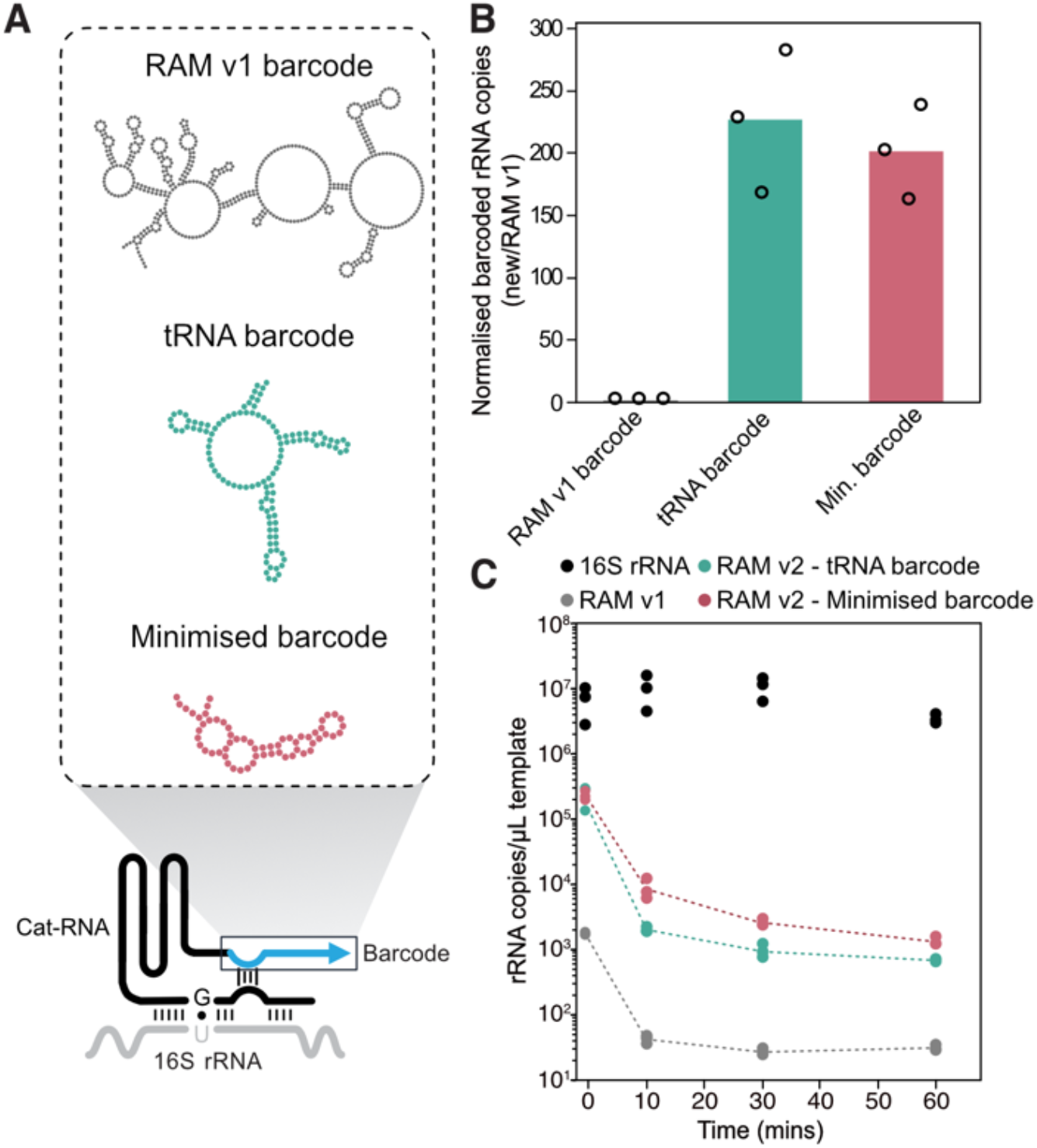
Barcode structure and length impact stability and signal of barcoded 16S rRNA. (**A**) Structures of the original barcode and the two rationally engineered variants. The first design incorporates a tRNA scaffold (tRNA barcode), while the second reduces sequence length while increasing secondary structure (minimised barcode). (**B**) Quantification of barcoded 16S rRNA using RT–qPCR in *E. coli* cells containing plasmids encoding RAM v1 (RAM v1 barcode) or the modified barcode designs, which include the P10 interaction. Barcoded 16S rRNA was normalised to native 16S rRNA. RT–qPCR data were normalised to the ‘RAM v1 barcode’ design. Barcoded rRNA signal for tRNA and minimised barcode designs is significantly higher than the RAM v1 barcode design (P<0.02 for both; *t-*test). (**C**) The stability of barcoded rRNA was compared with native rRNA for the three barcode designs. RT-qPCR was performed on *E. coli* cells containing plasmids encoding the three designs at different time points after rifampicin addition, which inhibits transcription. Individual data points show three biological replicates, and bars show the mean.

We next evaluated the stability of the barcoded rRNA products by quantifying their degradation rates. To do this, we arrested cellular transcription using a rifampicin spike-in (32) and extracted total RNA at 0, 10, 30, and 60 minutes following arrest. Native and barcoded 16S rRNA were quantified using RT-qPCR. From this, we saw that not only was the initial concentration (t=0) of RAM v1 lower than the other barcode designs, it also degraded rapidly and reached its minimal level after 30 minutes while the tRNA and minimised barcodes continued to drop until 60 minutes (Fig. 3C). We also confirmed that the new designs with increased 16S rRNA barcoding did not result in measurable growth rate defects in either rich or minimal media (Supplementary Fig. 6). Taken together, these results show the RAM signal can be substantially improved by more than 200-fold through a combination of rational ribozyme and barcode engineering. To distinguish these new RAM reporter designs, we herein refer to them as RAM version 2 (RAM v2).

### Modular RAM allows for predictable exchange of promoter elements

A practical consideration when using the RAM reporter to measure horizontal gene transfer by conjugation is that RAM transcription must be repressed in donor cells to avoid overwhelming the signal from transconjugants. To achieve this, we previously used a donor containing a non-transferrable plasmid that constitutively expresses the transcription factor CymR and placed the RAM reporter under the control of the corresponding promoter, Pcym (Fig. 4A). This configuration enabled ∼86-fold repression of RAM v1 transcription in the donor cell (18). However, as the CymR operator sites overlap with the promoter element, this repression mechanism is inherently restricted to the Pcym promoter. As a result, the promoter driving RAM cannot be readily exchanged. This constraint limits flexibility in modifying promoter strength, expanding the host range using broad-host-range promoters (19), exchanging donor strains, or enhancing RAM transcription in specific taxonomic groups (e.g., Gram-positive bacteria) (21). Additionally, one CymR operator is located downstream of the transcription start site (TSS), meaning this sequence is amended onto the cat-RNA, where it may interfere with function by forming intramolecular interactions.

**Figure 4.**
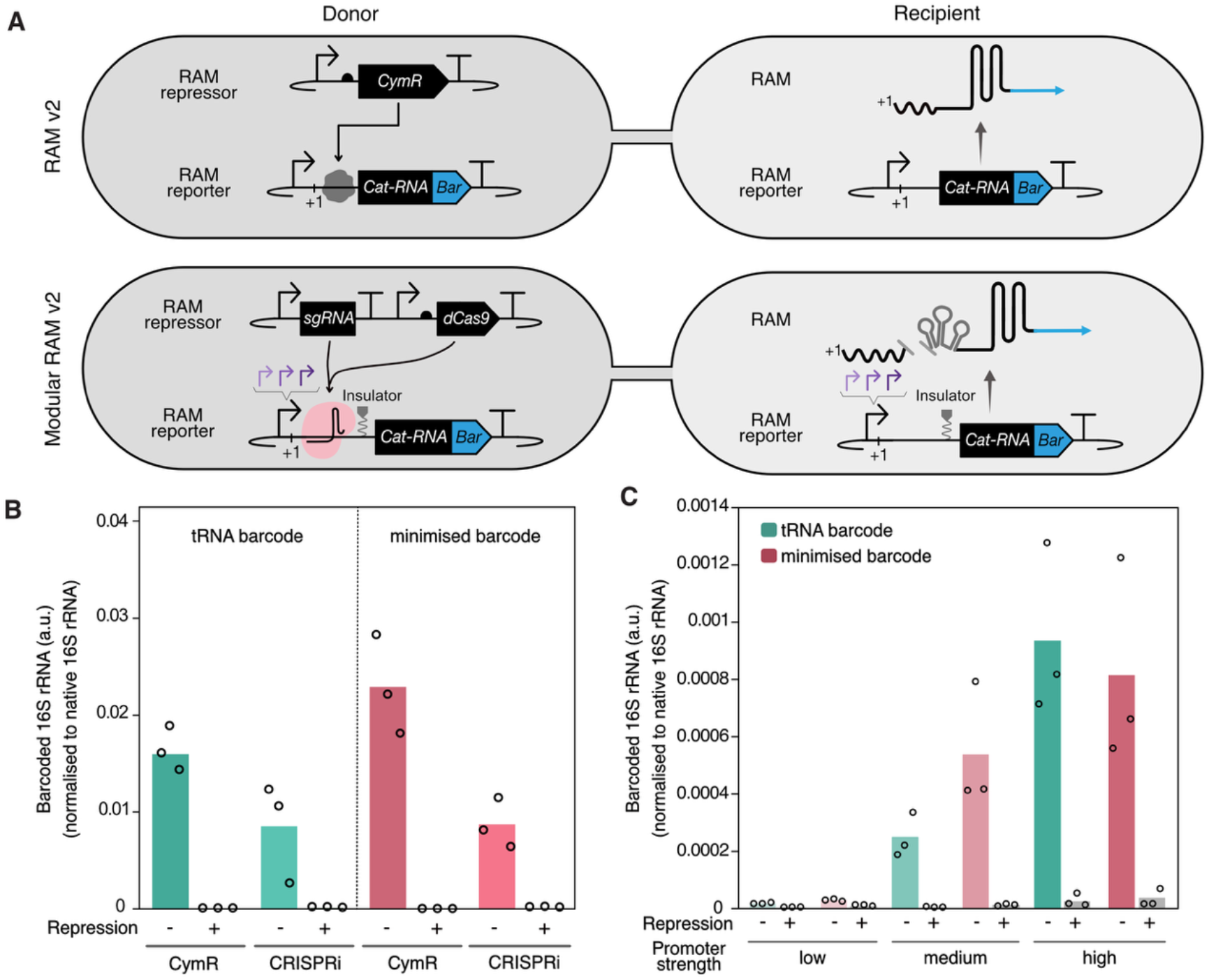
Modular RAM allows for predictable exchange of promoter elements. (**A**) (Top panel) Schematic of non-modular genetic design used for RAM v2, which uses the CymR promoter (Pcym) to drive RAM transcription, which is repressible in the donor by a plasmid encoding transcription factor CymR. (Bottom panel) In the modular genetic design, CRISPRi is used to repress the promoter in the donor, enabling facile promoter swapping. Additionally, a PlmJ insulator is included to cleave any amended sequences at the 5’ end of RAM. (**B**) Quantification of barcoded 16S rRNA using RT-qPCR in *E. coli* cells containing plasmids encoding either non-modular or modular RAM v2, with and without the corresponding repression system present (CymR or CRISPRi). Both constructs use Pcym to drive transcription of the RAM reporter. (**C**) Quantification of barcoded 16S rRNA using RT-qPCR in *E. coli* cells containing plasmids encoding the modular design. Three promoters of different strengths (J23105, J23101 and J23119) replaced the original promoter. Individual data points show three biological replicates, and bars represent the mean.

To address this, we designed a modular genetic construct for RAM v2. First, to establish a more generalisable repression mechanism, we used a CRISPR interference (CRISPRi) system that uses a catalytically dead Cas9 protein (dCas9) targeted to block transcription (26). CRISPRi has been reported to have high repression efficiency and a broad host range (33), making it well-suited for our application. To implement this design, a guide RNA (gRNA) recognition site was placed downstream of the promoter in the RAM cassette, and the corresponding gRNA and dCas9 were encoded on a separate non-transferrable plasmid. This makes it generalisable as the repression is independent of the promoter. To prevent sequences after the TSS from interfering with RAM, we incorporated the PlmJ insulator (27), which cleaves the upstream transcript, yielding a consistent cat-RNA transcript regardless of the upstream transcript (Fig 4A).

To evaluate the efficiency of the modular CRISPRi-based repression system, we compared it to the original CymR system (herein referred to as the non-modular design) by placing both constructs downstream of the same Pcym promoter. Cells were transformed with plasmids encoding either the non-modular or modular RAM v2, in the presence or absence of a plasmid encoding the corresponding repression system (CymR or CRISPRi). We evaluated both the tRNA and minimised barcode designs. Barcoded 16S rRNA levels were quantified, revealing strong RAM signal in the absence of repression and minimal signal in its presence for both the modular and non-modular designs (Fig. 4B, Supplementary Fig. 7). Interestingly, the modular design exhibited a modest reduction in signal under no-repression conditions, which we subsequently confirmed was due to the presence of the PlmJ insulator (Supplementary Fig. 8). Overall, these results demonstrate that the modular design achieves comparable performance to the original design while enabling promoter flexibility.

To further validate modularity, we replaced Pcym with three constitutive promoters of varying strengths: weak (J23105), medium (J23101) and strong (J23119). We quantified the concentration of barcoded 16S rRNA for each construct and the amount of barcoded 16S rRNA in cells containing the repression plasmid (Fig. 4C, Supplementary Fig. 9). As predicted, we observed varying levels of barcoded rRNA corresponding to promoter strength in the absence of dCas9. In the presence of the CRISPRi repression plasmid, we observe significant repression across all constructs. Taken together, these results demonstrate that modular RAM v2 enables predictable control of RAM expression, facilitates facile promoter swapping, and enhances its versatility as a reporter for horizontal gene transfer in microbial communities.

### RAM v2 improves the signal of gene transfer measurements in a complex microbial community

To validate that RAM v2 functions robustly in microbial communities, we measured its ability to detect conjugation of a plasmid from an *E. coli* donor into a wastewater microbial community. To benchmark performance, we directly compared RAM v1 and RAM v2. Two separate donor cells were prepared, each carrying a pBBR1 mobilisable plasmid encoding either RAM v1 or RAM v2 with the tRNA barcode (Fig. 5A). To facilitate a direct comparison, both designs used the Pcym promoter to ensure comparable transcription levels. These donor cells were added in separate mating assays to the same community, which was derived from a wastewater-activated sludge sample collected from the West University Wastewater Treatment Plant in Houston, Texas. Following overnight conjugation, total RNA was extracted, and RT-qPCR was performed to quantify native and barcoded 16S rRNA. Consistent with our findings in *E. coli*, we observed over a 150-fold increase in signal in RAM v2 compared to RAM v1 (Fig. 5B, Supplementary Fig. 10). This confirms that the improved RAM v2 translates to robust performance within a complex microbial community.

**Figure 5.**
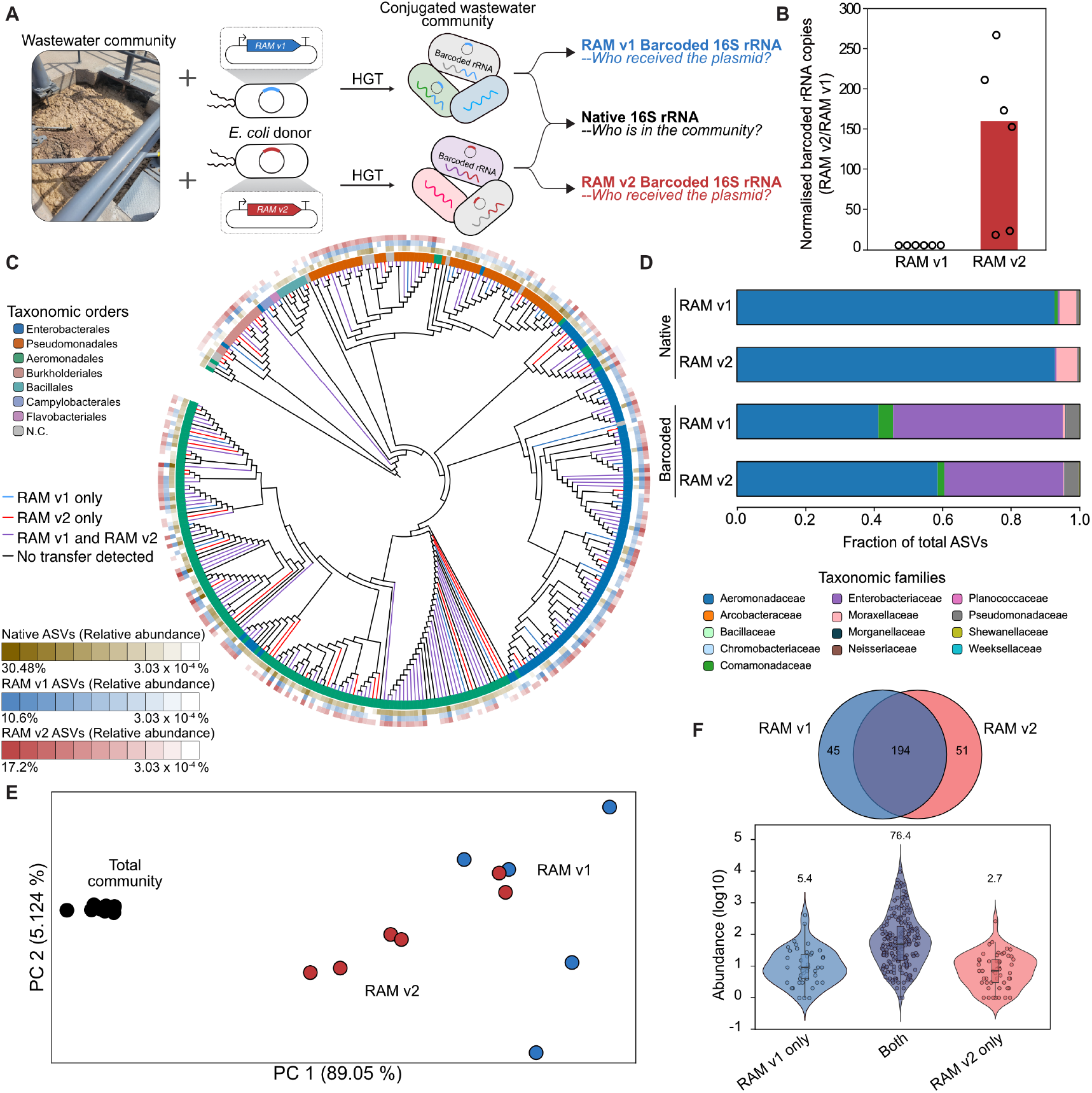
Comparing RAM designs for measuring conjugation in microbial communities. (**A**) RAM v1 and v2 were used to record information about conjugation within a Houston wastewater microbial community, and amplicon sequencing of the native and barcoded 16S rRNA showed which microorganisms were present and which microorganisms participated in conjugation. (**B**) Quantification of barcoded 16S rRNA using RT–qPCR in a diverse Houston wastewater microbial community conjugated with plasmids encoding RAM v1 or RAM v2 with the tRNA design. Barcoded 16S rRNA was normalised to native 16S rRNA. RT-qPCR data was normalised to RAM v1. Individual data points show three biological replicates, and bars represent the mean. Barcoded rRNA signal of RAM v2 was significantly higher than RAM v1 (P<0.02; *t-*test). (**C**) Conjugation host range mapped onto an evolutionary tree including all ASVs observed in the community. Blue branches represent ASVs barcoded by RAM v1, red branches represent ASVs barcoded by RAM v2, purple branches represent ASVs barcoded by both RAM v1 and v2, and black branches represent ASVs that were not barcoded. The relative abundances of native (brass), RAM v1 barcoded (blue) and RAM v2 barcoded (red) ASVs are shown, with the outer leaves showing taxonomic order. (**D**) The relative fraction of total ASVs observed in 14 different families. Each bar graph was generated from data collected across five biological replicates for RAM v1 and six for RAM v2. (**E**) PCoA reveals distinct clustering of RAM v1 and RAM v2 barcoded communities, which are cluster separately from the native community. PCoA was performed on the weighted unifrac distance matrix between populations. The significance of these differences was evaluated using permutational multivariate analysis of variance (in Supplementary figure 12). (**F**) Venn diagram of ASVs observed in RAM v1 and RAM v2, with the overlap representing ASVs observed in both. The corresponding violin plot shows the abundance of ASVs observed only in RAM v1, only in RAM v2, and both, with the average abundances for each category indicated above.

To characterise the community composition and identify transconjugants, amplicon sequencing of native and barcoded 16S rRNA was performed (Fig. 5C). A total of 345 amplicon sequence variants (ASVs) were detected across the native and transconjugant communities, spanning seven orders. Comparison of the native 16S rRNA profiles between RAM v1 and v2 mating assays showed that the native communities were highly similar, as expected (Fig. 5D). Among these ASVs, 239 and 245 were barcoded by RAM v1 and RAM v2, respectively (Supplementary Fig. 11). Across both designs, high levels of barcoding were observed for Aeromonadaceae, the most abundant family in the community (40% of ASVs), and for Enterobacteriaceae (20% of ASVs) (Fig. 5D).

To determine whether RAM v1 and v2 differed in the specific ASVs barcoded, we assessed compositional differences across samples using principal coordinate analysis (PCoA) (Fig. 5E). This analysis confirmed that the native communities in the RAM v1 and v2 mating assays were not significantly different. Interestingly, when comparing the RAM v1 and v2 barcoded 16S rRNA, we observed a significant difference in the transconjugant communities (Supplementary Fig. 12) (P=0.019; PERMANOVA). To identify ASVs driving the observed differences between the transconjugant communities, we looked at the unique or shared ASVs between RAM v1 and v2 (Fig. 5E). Of the ∼250 ASVs observed in the transconjugant communities, ∼200 were shared and ∼50 were unique to each version of RAM. Interestingly, we observed that the unique ASVs had lower average relative abundances than the shared ASVs (Fig. 5F). We also compared the relative abundances of the shared ASVs across RAM v1 and v2, which showed a strong correlation (R^2^ = 0.81) (Supplementary Fig. 13). These analyses suggest that rare events are driving the differences and may reflect the natural stochasticity of gene transfer among low-abundance community members. Overall, RAM v2 barcodes the same dominant taxa as RAM v1 and reveals a similar plasmid host range, proving to be a robust, reliable, and sensitive barcoding technology for studying plasmid host range.

## DISCUSSION

Our results demonstrate that rational design and engineering improved the sensitivity of an RNA barcoding system that can be used to record gene transfer in microbiomes. By engineering tertiary interactions into the ribozyme and a structured barcode that avoids cellular degradation, we established a RAM v2 reporter that shows hundreds of fold increase in signal in *E. coli* pure culture and a wastewater microbial community compared to RAM v1. Additionally, we established a modular RAM genetic architecture that enables facile and predictable promoter exchange. Taken together, these results deliver an improved RAM reporter that addresses fundamental bottlenecks in sensitivity, host-range constraints, and flexibility in expression levels within a microbial community.

Our work emphasises the value of using rational design alongside computational RNA structure prediction to deliver a RAM technology with stable and improved signals in complex microbial environments across multiple taxa. This work complements prior efforts to improve the function of synthetic RNA systems through structure-guided rational design (34–38). Beyond rational design, we anticipate that other strategies could be applied to get further gains in RAM functionality. For example, directed evolution (39) and high-throughput screening of mutant libraries have been widely applied to RNA systems (40), and these approaches could be readily adapted to enhance RAM activity. Beyond synthetic variants, the natural diversity of group I intron ribozymes could be explored to uncover new ribozymes with enhanced activity or distinct properties, such as smaller genetic footprints or enhanced specificity. Finally, several recent studies have highlighted the potential of using artificial intelligence for RNA design (41–43). Although these have focused on small RNAs, with continued advances in methods and bigger training datasets, the design of large, structurally complex RNAs will likely be possible in the coming years.

We anticipate that the enhanced efficiency of RAM v2 represents a major advance with significant implications for studying HGT within complex microbial communities. First, by improving splicing efficiency and reporter signal, RAM v2 will offer a lower limit of detection and thus has the potential to provide a higher-resolution picture of gene transfer by enabling the detection of rare events. Combining this enhanced sensitivity alongside RAM designed to specifically target rare taxa to enrich their signal within a community (44) would provide even greater enhancement. This is particularly powerful given that even low-abundance members of a biosphere can serve as primary drivers of genetic diversity (45). Likewise, this improved signal could facilitate the study of gene transfer in challenging environmental matrices where RNA isolation yields are typically low, such as soils (46). Secondly, the improved barcode design enables scalable experimental workflows, such as multiplexing RAM to record multiple gene transfer events in parallel. For example, the tRNA scaffold, shown to be a robust scaffold that allows expression and folding of diverse RNA sequences, could be leveraged to encode distinct barcode sequences. This would allow massive multiplexing of RAM measurements across diverse mobile genetic elements, including plasmids and bacteriophages.

Modular genetic design has greatly streamlined the exchange of genetic elements to optimise genetically encoded functions (47, 48). Herein, we present a RAM genetic construct with a modular architecture that facilitates the facile swapping of promoter elements. We anticipate this approach will be impactful in several ways. First, since promoters have distinct host ranges (49), tailoring these elements can enhance the transcription of the RAM reporter in specific taxonomic groups. Alternatively, the modular genetic design could be utilised to pair promoters and unique barcode sequences to allow for massively multiplexed promoter characterisation. Similarly, by utilising a CRISPRi repressor that has previously been shown to function across diverse microbes (50), our work allows the use of varied donor species in conjugation experiments. This allows the systematic exploration of donor-specific factors, such as surface exclusion (51), taxonomic relatedness (52), or restriction-modification systems (53), on gene transfer frequency and host range, parameters previously identified as critical. Finally, the RAM reporter provides a standardised system for delivering, tracking, and evaluating synthetic gene circuits in non-model microbes within microbial communities.

## METHODS

### Plasmid construction

Supplementary Table 1 lists all plasmids used in this study. Plasmids were constructed using PCR and Gibson assembly. Sanger sequencing or whole plasmid sequencing was used to verify the constructs. The tRNA barcode sequence was derived from *E. coli* tRNA-fMet. All RAM cassettes contain the pBBR1 origin of replication, and the repression plasmids contain the p15A origin of replication. The sequences of these origins of replication are provided as supporting information in Supplementary Table 2.

### Strains, growth and transformations

Supplementary Table 2 lists all strains used in this study. For all experiments, single colonies were used to inoculate liquid cultures in LB media with antibiotics. *E. coli* cells were transformed with plasmids using chemical transformation; all RAM plasmids containing kanamycin resistance cassettes were selected at a concentration of 100 µg/mL, and repression plasmids containing chloramphenicol resistance cassettes were selected at a concentration of 34 µg/mL.

### Growth measurements

Single colonies of *E. coli* cells transformed with plasmids were used to inoculate 500 µL LB media with corresponding antibiotics (LB-ab). After growing cultures overnight at 37 °C, shaking at 1000 rpm, 2 µL of culture was used to inoculate 198 µL fresh LB-ab medium in 96-well blocks. After 4 hours of growth, 1 µL was used to inoculate 99 µL of either LB or M9 medium supplemented with casamino acids and thiamine, with antibiotics in Nunc Edge 2.0 96-well plates (Thermo Fisher Scientific). Cells were grown for 17 h at 30 °C with double orbital rotation (amplitude: 3 mm) in a Tecan M1000 while measuring the optical density at 600 nm (OD600) every 10 min.

### Total RNA extraction

Single colonies of *E. coli* cells transformed with plasmids were used to inoculate 3 mL LB-ab medium and grown. After growing cultures at 37 °C shaking at 225 rpm, stationary-phase cultures were diluted 1:100 by adding 50 µL of culture in 5 mL LB-ab and grown for ∼4 h at 37 °C and 225 rpm. 500 µL of cells were pelleted by centrifugation (6,000 g, 5 min), and the Maxwell Purefood GMO and authentication kit (Promega) was used to extract total nucleic acids using an automatic magnetic bead purification method with a Maxwell RSC 48 instrument. In brief, cell pellets were resuspended by vortexing in 350 µL of CTAB buffer and 20 µL of Proteinase K, and incubated at 60 °C for 10 min. 300 µL of lysed sample and 300 µL of lysis buffer were added to the receiver well of the extraction cartridge, and nucleic acids were eluted in 50 µL of elution buffer. DNA was removed using a Turbo DNA-free kit (Invitrogen) by adding 2 µL of Turbo DNase to 20 µL of the eluate and incubating at 37 °C for 1 h. DNase was inactivated by adding 5 µL of the chelating bead suspension and agitating for 10 min at 2 min intervals to resuspend the beads. All samples were diluted tenfold to prevent the kit’s inhibitors from interfering with downstream reactions.

### Reverse transcription quantitative PCR (RT-qPCR)

Primers and probes used for RT-qPCR are described in Supplementary Table 4. RT-qPCR was performed using a dye-based qPCRbio SyGreen one-step kit (PCR Biosystems, PB25.11) in a QuantStudio-3 qPCR thermocycler (Thermo Fisher Scientific). Reactions were composed of 4 µL of RNA template, 0.4 µM of each primer, and 0.5 µL of the provided RTase Go reverse transcriptase in a 10 µL reaction volume. To compare the signal of barcoded 16S rRNA in a wastewater community conjugated with RAM v1 or RAM v2, RT-qPCRs were performed with probes using a PCRbio Probe one-step detect master mix (PCR Biosystems, PB25.41). Reactions were composed of 4 µL of RNA template, 0.4 µM of each primer, 0.2 µM of the probe and 0.1 µL of the provided RTase Go reverse transcriptase in a 10 µL reaction volume. Sample concentrations were converted to copies per µL of template using standard curves made with known concentrations of purified PCR products. Primer efficiencies were measured and are all within the 90-110% range. Melting curve analysis was performed, and a single peak was observed for all templates and products analysed using agarose gel electrophoresis, which confirmed the amplification of a single product.

### RNA stability measurements

To measure RNA stability, *E. coli* strains expressing different RAM constructs (RAM v1, RAM v2–tRNA or RAM v2-minimised) were grown overnight in 5 mL LB-ab media. 200 µL of this culture was added to 10 mL of LB-ab and grown for ∼3 hours. These cultures were then adjusted to an OD600 of 0.5 in fresh LB-ab containing rifampicin (150 ng/µL) to halt transcription. 750 µL cells were collected at the 0, 10, 30, and 60 min timepoints after incubation and frozen at -80 °C before RNA extraction and RT-qPCR.

### Wastewater conjugation and RNA extraction

Fifty mL of mixed liquor was collected on 15^th^ October 2025 from the aeration basin of the West University Wastewater Treatment plant in Houston, Texas. The sample was placed on ice and transported back to the laboratory, and conjugation was performed within an hour of sample collection. *E. coli* MFDpir was used as the conjugation donor. Donor cells containing plasmids that express CymR and RAM v1 or RAM v2 – tRNA were grown overnight in 1 mL LB-ab media containing 0.3 mM diaminopimelic acid (DAP). After growth, 1 mL of 1 OD_600_ cells was collected and washed three times with PBS to remove any antibiotics. In parallel, 1 mL of the wastewater sample was centrifuged at 13000 rcf for 3 min, and the pellet was resuspended in 1 mL of PBS. The donor cells and wastewater microbial community were mixed in a 1:1 v/v ratio, and 50 µL of each mixture was spotted onto a nitrocellulose filter placed on an LB-agar plate containing 0.3 mM DAP. After incubating at room temperature (25°C) for 24 hours, the filters were placed in a 2 mL tube, and the cells were resuspended in 500 µL of Cetyltrimethylammonium bromide (CTAB) buffer by vortexing. The filter paper was removed, and the suspension was heated at 95 °C for 5 min. Samples were cooled to room temperature and vortexed. 20 µL of proteinase K was added to each sample, and the samples were heated at 70 °C for 10 min. The extraction cartridge was prepared as explained in the above-mentioned RT-qPCR method. RNA obtained after DNase treatment was stored at -80 °C.

### Amplicon sequencing of the wastewater microbial community native and barcoded 16S rRNA

To convert extracted RNA to cDNA for amplicon sequencing, 0.5 µL of 2 µM RT primer, 0.5 µL of 10 mM dNTPs, ∼800 ng of purified RNA and water were combined for a final volume of 6.75 µL. Blocking oligonucleotides at a final concentration of 0.25 µM were used during conversion of the barcoded 16S rRNA to cDNA to maximise the production of correctly spliced products. RNA was denatured at 65 °C for 5 min and then cooled on ice. The following reagents (Thermo Fisher Scientific) were then added: 2 µL of 5x Superscript IV buffer, 0.5 µL of 100 mM dithiothreitol, 0.5 µL of 40 U/µL RNaseOUT ribonuclease inhibitor, and 0.25 µL of Superscript IV reverse transcriptase. RT was performed at 55 °C for 20 min and heat-inactivated at 80 °C for 10 min. The resulting cDNA was stored at -80 °C.

To amplify the native 16S rRNA, we used the universal 968F and 1391R primers, and to amplify the barcoded 16S rRNA, we used 968F and a custom primer (LNK27A) that binds to the barcode. PCR was performed using 5 µL of 5x Q5 reaction buffer, 1.25 µL of 10 µM forward and reverse primers each, 0.5 µL of 10 mM dNTPs, 0.25 µL of Q5 polymerase, 2.5 µL of cDNA and nuclease-free water for a final volume of 25 µL. Additionally, 0.25 µL of 10 µM blocking oligonucleotides was added to the barcoded 16S rRNA PCRs. The following thermocycler protocol was used: 98 °C for 2 min, followed by 25 cycles of 98 °C for 30 s, 60 °C for 45 s, and 72 °C for 1 min. For barcode PCRs, this step was repeated for 5 more cycles for a total of 30 cycles. All products underwent a final extension at 72 °C for 2 min and cooled to 4 °C for 15 min. PCR products were run on a 2% agarose gel; the expected products were gel-purified and used as a template for the second PCR to add the partial Illumina adaptor sequence. For this PCR, 30 µL reactions were set up. 1.5 µL of the gel-extracted product was used as a template for native 16S rRNA amplification, and 10 µL of gel-extracted product was used as a template for the barcoded 16S rRNA amplification. 6 µL of 5x Q5 reaction buffer, 1.5 µL of 10 µM forward and reverse primers, template, 0.3 µL of Q5 polymerase and nuclease-free water for a final volume of 30 µL were combined. The thermocycler settings mentioned above were used for native and barcoded 16S rRNA amplification, except for the number of cycles - native 16S rRNA underwent 10 amplification cycles, and the barcoded 16S rRNA underwent 12 cycles. 2 µL of the product was run on a gel to verify target amplification, and the remaining products were column-purified. These products were sequenced using the Amplicon-EZ service by Genewiz. All primers and blocking oligonucleotides are shown in Supplementary Table 4.

### Data analysis pipeline

The NGS reads were imported into QIIME 2 (version 2024.10) (54), and Cutadapt was used to trim the raw reads so that both the native and barcoded 16S rRNA sequences contained the same homologous regions. To do this, the forward and reverse primers (AACGCGAAGAACCTTAC and CGTTTGACGGGCGGTGWGTRCA) were trimmed from the native reads, and a short sequence after the splice site (AGGCCCGGGAACGT) was also trimmed to match the amplicons from the barcoded reads. To trim the barcoded reads, the forward and reverse primers, along with a short 2-nucleotide sequence (AACGCGAAGAACCTTAC and GGGAAAAGCATTGAACACCAT), were removed. The resulting sequences across the native and barcoded reads were now completely homologous. The DADA2 plugin was used to filter, denoise and dereplicate the reads. This provided a feature table of each unique ASV, its relative abundance, and the total counts for each ASV per sample. This feature table was filtered to retain ASVs with a total sequence length of ≥ 250 nt and had at least three total reads, and all samples were rarefied to 15000 reads. Taxonomy was assigned to each ASV using `feature-classifier classify-sklearn`, and a naive Bayes classifier (silva-138-99) was trained on the amplified 16S rRNA region. The plugin `fragment-insertion sepp` was used to generate a phylogenetic tree of filtered ASVs using a fragment-insertion approach. Interactive Tree of Life (iTOL) (55) was used to annotate the phylogenetic tree. The weighted unifrac distance matrix and corresponding PCoA plot were calculated using `diversity-lib weighted-unifrac` and `diversity beta-group-significance` plugins.

### Statistical analysis

A Welch *t*-test was used to compare variants to RAM v1 in all experiments. P < 0.05 (two-tailed) was considered for statistical significance.

## ACKNOWLEDGEMENTS

We thank Drs. Prashant B. Kalvapalle and August Staubus for their valuable input and discussions on ribozyme design and data analysis, and Darren Seet and Malyn Selinidis for their protocols and help with data analysis. We also thank the West University Wastewater Treatment Plant operators for their assistance with collecting samples. This work was supported by the following funding agencies: National Science Foundation (CAREER grant no. 2237052 to L.B.S.; CAREER grant no. 2237512 to J.C.; grant no. 2516327 to L.B.S. and J.C.) and the Army Research Office (W911NF-24-2-0073 to J.J.S., J.C. and L.B.S.). The views and conclusions contained in this document are those of the authors and should not be interpreted as representing the official policies, either expressed or implied, of the Army Research Office or the US Government. The US Government is authorised to reproduce and distribute reprints for Government purposes, notwithstanding any copyright notation.

