## Supplementary Fig. for "Rational engineering enhances the signal and modularity of an RNA barcoding technology to track gene transfer in microbiomes"



| | |
|---|---|
| **Pages 2-4** | Supplementary table 1. Plasmids used in this study. |
| **Pages 4-8** | Supplementary table 2. Key plasmid DNA and barcoded 16S rRNA sequences. |
| **Page 8** | Supplementary table 3. Strains used in this study. |
| **Pages 9** | Supplementary table 4. Primers used for RT-qPCR and NGS. |
| **Page 10** | Supplementary Figure 1. A schematic of the wild-type group I intron splicing ribozyme from *Tetrahymena thermophila*. |
| **Page 11** | Supplementary Figure 2. P10 interaction improves splicing efficiency. |
| **Page 12** | Supplementary Figure 3. Schematic of different RAM barcode designs. |
| **Page 13** | Supplementary Figure 4. Barcode structure and length, combined with P10 interaction, increase barcoded 16S rRNA signal. |
| **Page 14** | Supplementary Figure 5. Barcode structure and length impact the signal of barcoded 16S rRNA. |
| **Page 15** | Supplementary Figure 6. Rationally engineered RAM does not affect *E. coli* cell fitness. |
| **Page 16** | Supplementary Figure 7: Repression of RAM and modular RAM in *E. coli* cells. |
| **Page 17** | Supplementary Figure 8. RiboJ in modular RAM v2 design results in a drop in barcoded 16S rRNA signal. |
| **Page 18** | Supplementary Figure 9. Modular RAM v2 works robustly across different promoter strengths. |
| **Page 19** | Supplementary Figure 10. RAM v2 has increased barcode signal in a diverse Houston wastewater microbial community. |
| **Page 20** | Supplementary Figure 11. Native, RAM v1 barcoded, and RAM v2 barcoded ASVs observed in the wastewater microbial community. |
| **Page 21** | Supplementary Figure 12. PERMANOVA of a weighted unifrac distance matrix between the native and barcoded ASVs of RAM v1 and v2. |
| **Page 22** | Supplementary Figure 13. Scatter plot of ASVs barcoded by RAM v1 and RAM v2. |

**Supplementary Table 1. Plasmids used in this study**. For each plasmid, we note the use, the relevant data shown, and the plasmid architecture and features, including: (i) selectable marker, (ii) origin of replication, (iii) promoter, (iv) translation initiation sequences, (v) open reading frames, e.g., mGreenLantern, and (vi) ribozyme barcode details.

| Plasmid name | Name | Use | Figure | Plasmid Architecture |
|---|---|---|---|---|
| pPK076 | RAM v1 | RAM v1 splicing variant | 2B, 3B, 3C, 5 | KanR - pBBR1 - PcymRC - U1376_Target_50nt – Ribozyme – Barcode (sfGFP_2) - tVoigtS4 |
| pLNK001 | WT P10 | P10 variants | 2B | KanR - pBBR1 - PcymRC - U1376_Target_50nt - WT P10 - Ribozyme – Barcode (sfGFP_2) - tVoigtS4 |
| pLNK002 | Syn. P10 | | 2B | KanR - pBBR1 - PcymRC - U1376_Target_50nt - Syn. P10 - Ribozyme – Barcode (sfGFP_2) - tVoigtS4 |
| pLNK003 | tRNA barcode | barcode variants | SI 5 | KanR - pBBR1 - PcymRC - U1376_Target_50nt - Ribozyme – tRNA barcode - tVoigtS4 |
| pLNK004 | min. barcode | | SI 5 | KanR - pBBR1 - PcymRC - U1376_Target_50nt - Ribozyme – minimised barcode - tVoigtS4 |
| pLNK007 | RAM v2 - tRNA barcode | RAM v2 - Syn. P10 + barcode variants | 3B, 4B, 5 | KanR - pBBR1 - PcymRC - U1376_Target_50nt - Syn. P10 - Ribozyme – tRNA barcode - tVoigtS4 |
| pLNK008 | RAM v2 - min. barcode | | 3B, 4B | KanR - pBBR1 - PcymRC - U1376_Target_50nt - Syn. P10 - Ribozyme – minimised barcode - tVoigtS4 |
| pLNK013 | modular RAM v2 - tRNA barcode | RAM v2 variants (dCas9 repressible) with modular architecture | 4B | KanR - pBBR1 - PcymRC - gRNA target - PlmJ - U1376_Target_50nt - Syn. P10 - Ribozyme – tRNA barcode - tVoigtS4 |
| pLNK014 | modular RAM v2 - min. barcode | | 4B | KanR - pBBR1 - PcymRC - gRNA target - PlmJ - U1376_Target_50nt - Syn. P10 - Ribozyme – minimised barcode - tVoigtS4 |
| pLNK015 | modular (No PlmJ) RAM v2 - tRNA barcode | RAM v2 (dCas9 repressible) variants | SI 8 | KanR - pBBR1 - PcymRC - gRNA target - U1376_Target_50nt - Syn. P10 - Ribozyme – tRNA barcode - tVoigtS4 |

| | | | | |
|---|---|---|---|---|
| pLNK016 | modular (No PlmJ) RAM v2 - min. barcode | without PlmJ insulator | SI 8 | KanR - pBBR1 - PcymRC - gRNA target - U1376_Target_50nt - Syn. P10 - Ribozyme – minimised barcode - tVoigtS4 |
| pLNK018 | low exp. RAM v2 - tRNA barcode | | 4C | KanR - pBBR1 - J23105 - gRNA target - PlmJ - U1376_Target_50nt - Syn. P10 - Ribozyme – tRNA barcode - tVoigtS4 |
| pLNK019 | low exp. RAM v2 - min. barcode | | 4C | KanR - pBBR1 - J23105 - gRNA target - PlmJ - U1376_Target_50nt - Syn. P10 - Ribozyme – minimised barcode - tVoigtS4 |
| pLNK020 | med. exp. RAM v2 - tRNA barcode | RAM v2 variants expressed using promoters of varying strengths | 4C | KanR - pBBR1 - J23101 - gRNA target - PlmJ - U1376_Target_50nt - Syn. P10 - Ribozyme – tRNA barcode - tVoigtS4 |
| pLNK021 | med. exp. RAM v2 - min. barcode | | 4C | KanR - pBBR1 - J23101 - gRNA target - PlmJ - U1376_Target_50nt - Syn. P10 - Ribozyme – minimised barcode - tVoigtS4 |
| pLNK022 | high exp. RAM v2 - tRNA barcode | | 4C | KanR - pBBR1 - J23119 - gRNA target - PlmJ - U1376_Target_50nt - Syn. P10 - Ribozyme – tRNA barcode - tVoigtS4 |
| pLNK023 | high exp. RAM v2 - min. barcode | | 4C | KanR - pBBR1 - J23119 - gRNA target - PlmJ - U1376_Target_50nt - Syn. P10 - Ribozyme – minimised barcode - tVoigtS4 |

| | | | | |
|---|---|---|---|---|
| pLNK017 | dCas9 for RAM repression | modular RAM repression using dCas9 and gRNA | 4B, 4C | p15A - CmR - pTetR - dCas9 - dblTerm - J23119 - sgRNA - sgRNA scaffold - T500 |
| pPK067 | CymR repressor | Suppressing ribozyme using CymR | 4B, 5 | CmR - p15A - PlacIq - CymR - RiboJ100 - mGreenLantern - tVoigtN1 |
| pJEC905 | empty control | Negative control for ribozyme | SI 6 | KanR – pBBR1 |

**Supplementary Table 2. Key plasmid DNA and barcoded 16S rRNA sequences.** Features are highlighted with colours corresponding to the DNA sequence.

| Name and features | DNA sequence |
|---|---|
| Plasmid RAM v1 (PcymRC-U1376_Target_50nt – Ribozyme – Barcode (sfGFP_2)-tVoigtS4) | AACAAACAGACAATCTGGTCTGTTTGTATTATGGAAAATTTTTCTGTATAATA<br>GATTCAACAAACAGACAATCTGGTCTGTTTGTATTATCGACCAACCCACTCC<br>CATGGTGTGACGGGCGGTGTGTACAAGGCCCGGGAACGTGTTCACAAAAG<br>TTATCAGGCATGCACCTGGTAGCTAGTCTTTAAACCAATAGATTGCATCGGT<br>TTAAAAGGCAAGACCGTCAAATTGCGGGAAAGGGGTCAACAGCCGTTCAG<br>TACCAAGTCTCAGGGGAAACTTTGAGATGGCCTTGCAAAGGGTATGGTAAT<br>AAGCTGACGGACATGGTCCTAACCACGCAGCCAAGTCCTAAGTCAACAGAT<br>CTTCTGTTGATATGGATGCAGTTCACAGACTAAATGTCGGTCGGGGAAGAT<br>GTATTCTTCTCATAAGATATAGTCGGACCTCTCCTTAATGGGAGCTAGCGG<br>ATGAAGTGATGCAACACTGGAGCCGCTGGGAACTAATTTGTATGCGAAAGT<br>ATATTGATTAGTTTTGGAGTACTCGATGGTGTTCAATGCTTTTCCCGTTATC<br>CGGATCACATGAAACGGCATGACTTTTTCAAGAGTGCCATGCCCGAAGGTT<br>ATGTACAGGAACGCACTATATCTTTCAAAGATGACGGGACCTACAAGACGC<br>GTGCTGAAGTCAAGTTTGAAGGTGATACCCTTGTTAATCGTATCGAGTTAAA<br>GGGTATTGATTTTAAAGAAGATGGAAACATTCTTGGACACAAACTCGAGTAC<br>AACTTTAACTCACACAATGTATACATCACGGCAGACAAACAAAAGAATGGAA<br>TCAAAGCTAACTTCAAAATTCGCCACAACGTTGAAGATGGTTCCGTTCAACT<br>AGCAGACCATTATCAACAAAATACTCCAATTGGCGATGGCCCTGTCCTTTTA<br>CCAGACAACCATTACCTGTCGACACAATCTGTCCTTTCGAAAGATCCCAAC<br>GAAAAGCGTGACCACATGGTCCTTCTTGAGTTTGTAACTGCTGCTGGGATT<br>ACACATGGCATGGATGAGCTCTACAAATGGCCCAATTATTGAAGGCCTCCC<br>TAACGGGGGGCCTTTTTTTGTTTCTGGTCTGCCGCTG |
| Plasmid Syn. P10 | AACAAACAGACAATCTGGTCTGTTTGTATTATGGAAAATTTTTCTGTATAATA<br>GATTCAACAAACAGACAATCTGGTCTGTTTGTATTATCGACCTTCTTTTGCA<br>ACCCACTCCCATGGTGTGACGGGCGGTGTGTACAAGGCCtAACCACAcgTA<br>AAAGTTATCAGGCATGCACCTGGTAGCTAGTCTTTAAACCAATAGATTGCAT |

| | |
|---|---|
| (PcymRC-U1376_Target_50nt – Syn. P10 - Ribozyme – Barcode (sfGFP_2)-tVoigtS4) | CGGTTTAAAAGGCAAGACCGTCAAATTGCGGGAAAGGGGTCAACAGCCGT<br>TCAGTACCAAGTCTCAGGGGAAACTTTGAGATGGCCTTGCAAAGGGTATGG<br>TAATAAGCTGACGGACATGGTCCTAACCACGCAGCCAAGTCCTAAGTCAAC<br>AGATCTTCTGTTGATATGGATGCAGTTCACAGACTAAATGTCGGTCGGGGA<br>AGATGTATTCTTCTCATAAGATATAGTCGGACCTCTCCTTAATGGGAGCTAG<br>CGGATGAAGTGATGCAACACTGGAGCCGCTGGGAACTAATTTGTATGCGAA<br>AGTATATTGATTAGTTTTGGAGTACTCGATGGTGTTCAATGCTTTTCCCGTT<br>ATCCGGATCACATGAAACGGCATGACTTTTTCAAGAGTGCCATGCCCGAAG<br>GTTATGTACAGGAACGCACTATATCTTTCAAAGATGACGGGACCTACAAGA<br>CGCGTGCTGAAGTCAAGTTTGAAGGTGATACCCTTGTTAATCGTATCGAGT<br>TAAAGGGTATTGATTTTAAAGAAGATGGAAACATTCTTGGACACAAACTCGA<br>GTACAACTTTAACTCACACAATGTATACATCACGGCAGACAAACAAAAGAAT<br>GGAATCAAAGCTAACTTCAAAATTCGCCACAACGTTGAAGATGGTTCCGTT<br>CAACTAGCAGACCATTATCAACAAAATACTCCAATTGGCGATGGCCCTGTC<br>CTTTTACCAGACAACCATTACCTGTCGACACAATCTGTCCTTTCGAAAGATC<br>CCAACGAAAAGCGTGACCACATGGTCCTTCTTGAGTTTGTAACTGCTGCTG<br>GGATTACACATGGCATGGATGAGCTCTACAAATGGCCCAATTATTGAAGGC<br>CTCCCTAACGGGGGGCCTTTTTTTGTTTCTGGTCTGCCGCTG |
| Plasmid RAM v2 – tRNA barcode (PcymRC-U1376_Target_50nt – Syn. P10 - Ribozyme – tRNA barcode - tVoigtS4) | AACAAACAGACAATCTGGTCTGTTTGTATTATGGAAAATTTTTCTGTATAAT<br>AGATTCAACAAACAGACAATCTGGTCTGTTTGTATTATCGACCTTCTTTTGC<br>AACCCACTCCCATGGTGTGACGGGCGGTGTGTACAAGGCCtAACCACAcgT<br>AAAAGTTATCAGGCATGCACCTGGTAGCTAGTCTTTAAACCAATAGATTGC<br>ATCGGTTTAAAAGGCAAGACCGTCAAATTGCGGGAAAGGGGTCAACAGCC<br>GTTCAGTACCAAGTCTCAGGGGAAACTTTGAGATGGCCTTGCAAAGGGTA<br>TGGTAATAAGCTGACGGACATGGTCCTAACCACGCAGCCAAGTCCTAAGT<br>CAACAGATCTTCTGTTGATATGGATGCAGTTCACAGACTAAATGTCGGTCG<br>GGGAAGATGTATTCTTCTCATAAGATATAGTCGGACCTCTCCTTAATGGGA<br>GCTAGCGGATGAAGTGATGCAACACTGGAGCCGCTGGGAACTAATTTGTA<br>TGCGAAAGTATATTGATTAGTTTTGGAGTACTCGATGGTGTTCAATGCTTTT<br>CCCCGCGGGGTGGAGCAGCCTGGTAGCTCGTCGGAGTGCAAAGATGACG<br>GGACCTACATCACCCGAAGATCGTCGGTTCAAATCCGGCCCCCGCAACCA<br>TGGCCCAATTATTGAAGGCCTCCCTAACGGGGGGCCTTTTTTTGTTTCTGG<br>TCTGCCGCTG |
| Plasmid Modular RAM v2 – minimised barcode (PcymRC-guide RNA target – PlmJ - U1376_Target_50nt – Syn. | AACAAACAGACAATCTGGTCTGTTTGTATTATGGAAAATTTTTCTGTATAAT<br>AGATTCAACAAACAGACAATCTGGTCTGTTTGTATTATCGACcctTGGAACC<br>GTACTGGAACTGCtggagtcataagtctgggctaagcccactgatgagtcgctgaaatgcgacgaa<br>acttatgacctctacaaataattttgtttaaCTTCTTTTGCAACCCACTCCCATGGTGTGAC<br>GGGCGGTGTGTACAAGGCCtAACCACAcgTAAAAGTTATCAGGCATGCACC<br>TGGTAGCTAGTCTTTAAACCAATAGATTGCATCGGTTTAAAAGGCAAGACC<br>GTCAAATTGCGGGAAAGGGGTCAACAGCCGTTCAGTACCAAGTCTCAGGG<br>GAAACTTTGAGATGGCCTTGCAAAGGGTATGGTAATAAGCTGACGGACAT<br>GGTCCTAACCACGCAGCCAAGTCCTAAGTCAACAGATCTTCTGTTGATATG<br>GATGCAGTTCACAGACTAAATGTCGGTCGGGGAAGATGTATTCTTCTCATA<br>AGATATAGTCGGACCTCTCCTTAATGGGAGCTAGCGGATGAAGTGATGCA |

| | |
|---|---|
| P10 - Ribozyme – minimised barcode - tVoigtS4) | ACACTGGAGCCGCTGGGAACTAATTTGTATGCGAAAGTATATTGATTAGTT<br>TTGGAGTACTCGATGGTGTTCAATGCTTTTCCCAGACGCGTGCTGAAGTC<br>AAGCAAAGATGACGGGACCTACATGGCCCAATTATTGAAGGCCTCCCTAA<br>CGGGGGGCCTTTTTTTGTTTCTGGTCTGCCGCTG |
| RAM v1 barcoded 16S rRNA (16S - gfp barcode) | AAATTGAAGAGTTTGATCATGGCTCAGATTGAACGCTGGCGGCAGGCCTA<br>ACACATGCAAGTCGAACGGTAACAGGAAGAAGCTTGCTTCTTTGCTGACG<br>AGTGGCGGACGGGTGAGTAATGTCTGGGAAACTGCCTGATGGAGGGGGA<br>TAACTACTGGAAACGGTAGCTAATACCGCATAACGTCGCAAGACCAAAGA<br>GGGGTACCTTCGGGCCTCTTGCCATCGGATGTGCCCAGATGGGATTAGCT<br>AGTAGGTGGGGTAACGGCTCACCTAGGCGACGATCCCTAGCTGGTCTGA<br>GAGGATGACCAGCCACACTGGAACTGAGACACGGTCCAGACTCCTACGG<br>GAGGCAGCAGTGGGGAATATTGCACAATGGGCGCAAGCCTGATGCAGCC<br>ATGCCGCGTGTATGAAGAAGGCCTTCGGGTTGTAAAGTACTTTCAGCGGG<br>GAGGAAGGGAGTAAAGTTAATACCTTTGCTCATTGACGTTACCCGCAGAA<br>GAAGCACCGGCTAACTCCGTGCCAGCAGCCGCGGTAATACGGAGGGTGC<br>AAGCGTTAATCGGAATTACTGGGCGTAAAGCGCACGCAGGCGGTTTGTTA<br>AGTCAGATGTGAAATCCCCGGGCTCAACCTGGGAACTGCATCTGATACTG<br>GCAAGCTTGAGTCTCGTAGAGGGGGGTAGAATTCCAGGTGTAGCGGTGA<br>AATGCGTAGAGATCTGGAGGAATACCGGTGGCGAAGGCGGCCCCCTGGA<br>CGAAGACTGACGCTCAGGTGCGAAAGCGTGGGGAGCAAACAGGATTAGA<br>TACCCTGGTAGTCCACGCCGTAAACGATGTCGACTTGGAGGTTGTGCCCT<br>TGAGGCGTGGCTTCCGGAGCTAACGCGTTAAGTCGACCGCCTGGGGAGT<br>ACGGCCGCAAGGTTAAAACTCAAATGAATTGACGGGGGCCCGCACAAGC<br>GGTGGAGCATGTGGTTTAATTCGATGCAACGCGAAGAACCTTACCTGGTC<br>TTGACATCCACGGAAGTTTTCAGAGATGAGAATGTGCCTTCGGGAACCGT<br>GAGACAGGTGCTGCATGGCTGTCGTCAGCTCGTGTTGTGAAATGTTGGGT<br>TAAGTCCCGCAACGAGCGCAACCCTTATCCTTTGTTGCCAGCGGTCCGGC<br>CGGGAACTCAAAGGAGACTGCCAGTGATAAACTGGAGGAAGGTGGGGAT<br>GACGTCAAGTCATCATGGCCCTTACGACCAGGGCTACACACGTGCTACAA<br>TGGCGCATACAAAGAGAAGCGACCTCGCGAGAGCAAGCGGACCTCATAA<br>AGTGCGTCGTAGTCCGGATTGGAGTCTGCAACTCGACTCCATGAAGTCGG<br>AATCGCTAGTAATCGTGGATCAGAATGCCACGGTGAATATGGTGTTCAATG<br>CTTTTCCCGTTATCCGGATCACATGAAACGGCATGACTTTTTCAAGAGTGC<br>CATGCCCGAAGGTTATGTACAGGAACGCACTATATCTTTCAAAGATGACG<br>GGACCTACAAGACGCGTGCTGAAGTCAAGTTTGAAGGTGATACCCTTGTT<br>AATCGTATCGAGTTAAAGGGTATTGATTTTAAAGAAGATGGAAACATTCTT<br>GGACACAAACTCGAGTACAACTTTAACTCACACAATGTATACATCACGGCA |

| | |
|---|---|
| | GACAAACAAAAGAATGGAATCAAAGCTAACTTCAAAATTCGCCACAACGTT<br>GAAGATGGTTCCGTTCAACTAGCAGACCATTATCAACAAAATACTCCAATT<br>GGCGATGGCCCTGTCCTTTTACCAGACAACCATTACCTGTCGACACAATCT<br>GTCCTTTCGAAAGATCCCAACGAAAAGCGTGACCACATGGTCCTTCTTGA<br>GTTTGTAACTGCTGCTGGGATTACACATGGCATGGATGAGCTCTACAAA |
| RAM v2 tRNA barcoded 16S rRNA (16S - tRNA barcode) | AAATTGAAGAGTTTGATCATGGCTCAGATTGAACGCTGGCGGCAGGCCTA<br>ACACATGCAAGTCGAACGGTAACAGGAAGAAGCTTGCTTCTTTGCTGACG<br>AGTGGCGGACGGGTGAGTAATGTCTGGGAAACTGCCTGATGGAGGGGGA<br>TAACTACTGGAAACGGTAGCTAATACCGCATAACGTCGCAAGACCAAAGA<br>GGGGTACCTTCGGGCCTCTTGCCATCGGATGTGCCCAGATGGGATTAGCT<br>AGTAGGTGGGGTAACGGCTCACCTAGGCGACGATCCCTAGCTGGTCTGA<br>GAGGATGACCAGCCACACTGGAACTGAGACACGGTCCAGACTCCTACGG<br>GAGGCAGCAGTGGGGAATATTGCACAATGGGCGCAAGCCTGATGCAGCC<br>ATGCCGCGTGTATGAAGAAGGCCTTCGGGTTGTAAAGTACTTTCAGCGGG<br>GAGGAAGGGAGTAAAGTTAATACCTTTGCTCATTGACGTTACCCGCAGAA<br>GAAGCACCGGCTAACTCCGTGCCAGCAGCCGCGGTAATACGGAGGGTGC<br>AAGCGTTAATCGGAATTACTGGGCGTAAAGCGCACGCAGGCGGTTTGTTA<br>AGTCAGATGTGAAATCCCCGGGCTCAACCTGGGAACTGCATCTGATACTG<br>GCAAGCTTGAGTCTCGTAGAGGGGGGTAGAATTCCAGGTGTAGCGGTGA<br>AATGCGTAGAGATCTGGAGGAATACCGGTGGCGAAGGCGGCCCCCTGGA<br>CGAAGACTGACGCTCAGGTGCGAAAGCGTGGGGAGCAAACAGGATTAGA<br>TACCCTGGTAGTCCACGCCGTAAACGATGTCGACTTGGAGGTTGTGCCCT<br>TGAGGCGTGGCTTCCGGAGCTAACGCGTTAAGTCGACCGCCTGGGGAGT<br>ACGGCCGCAAGGTTAAAACTCAAATGAATTGACGGGGGCCCGCACAAGC<br>GGTGGAGCATGTGGTTTAATTCGATGCAACGCGAAGAACCTTACCTGGTC<br>TTGACATCCACGGAAGTTTTCAGAGATGAGAATGTGCCTTCGGGAACCGT<br>GAGACAGGTGCTGCATGGCTGTCGTCAGCTCGTGTTGTGAAATGTTGGGT<br>TAAGTCCCGCAACGAGCGCAACCCTTATCCTTTGTTGCCAGCGGTCCGGC<br>CGGGAACTCAAAGGAGACTGCCAGTGATAAACTGGAGGAAGGTGGGGAT<br>GACGTCAAGTCATCATGGCCCTTACGACCAGGGCTACACACGTGCTACAA<br>TGGCGCATACAAAGAGAAGCGACCTCGCGAGAGCAAGCGGACCTCATAA<br>AGTGCGTCGTAGTCCGGATTGGAGTCTGCAACTCGACTCCATGAAGTCGG<br>AATCGCTAGTAATCGTGGATCAGAATGCCACGGTGAATATGGTGTTCAATG<br>CTTTTCCCCGCGGGGTGGAGCAGCCTGGTAGCTCGTCGGAGTGCAAAGA<br>TGACGGGACCTACATCACCCGAAGATCGTCGGTTCAAATCCGGCCCCCG<br>CAACCA |

| | |
|---|---|
| RAM v2 minimised barcoded 16S rRNA (16S - minimised barcode) | AAATTGAAGAGTTTGATCATGGCTCAGATTGAACGCTGGCGGCAGGCCTA<br>ACACATGCAAGTCGAACGGTAACAGGAAGAAGCTTGCTTCTTTGCTGACG<br>AGTGGCGGACGGGTGAGTAATGTCTGGGAAACTGCCTGATGGAGGGGGA<br>TAACTACTGGAAACGGTAGCTAATACCGCATAACGTCGCAAGACCAAAGA<br>GGGGTACCTTCGGGCCTCTTGCCATCGGATGTGCCCAGATGGGATTAGCT<br>AGTAGGTGGGGTAACGGCTCACCTAGGCGACGATCCCTAGCTGGTCTGA<br>GAGGATGACCAGCCACACTGGAACTGAGACACGGTCCAGACTCCTACGG<br>GAGGCAGCAGTGGGGAATATTGCACAATGGGCGCAAGCCTGATGCAGCC<br>ATGCCGCGTGTATGAAGAAGGCCTTCGGGTTGTAAAGTACTTTCAGCGGG<br>GAGGAAGGGAGTAAAGTTAATACCTTTGCTCATTGACGTTACCCGCAGAA<br>GAAGCACCGGCTAACTCCGTGCCAGCAGCCGCGGTAATACGGAGGGTGC<br>AAGCGTTAATCGGAATTACTGGGCGTAAAGCGCACGCAGGCGGTTTGTTA<br>AGTCAGATGTGAAATCCCCGGGCTCAACCTGGGAACTGCATCTGATACTG<br>GCAAGCTTGAGTCTCGTAGAGGGGGGTAGAATTCCAGGTGTAGCGGTGA<br>AATGCGTAGAGATCTGGAGGAATACCGGTGGCGAAGGCGGCCCCCTGGA<br>CGAAGACTGACGCTCAGGTGCGAAAGCGTGGGGAGCAAACAGGATTAGA<br>TACCCTGGTAGTCCACGCCGTAAACGATGTCGACTTGGAGGTTGTGCCCT<br>TGAGGCGTGGCTTCCGGAGCTAACGCGTTAAGTCGACCGCCTGGGGAGT<br>ACGGCCGCAAGGTTAAAACTCAAATGAATTGACGGGGGCCCGCACAAGC<br>GGTGGAGCATGTGGTTTAATTCGATGCAACGCGAAGAACCTTACCTGGTC<br>TTGACATCCACGGAAGTTTTCAGAGATGAGAATGTGCCTTCGGGAACCGT<br>GAGACAGGTGCTGCATGGCTGTCGTCAGCTCGTGTTGTGAAATGTTGGGT<br>TAAGTCCCGCAACGAGCGCAACCCTTATCCTTTGTTGCCAGCGGTCCGGC<br>CGGGAACTCAAAGGAGACTGCCAGTGATAAACTGGAGGAAGGTGGGGAT<br>GACGTCAAGTCATCATGGCCCTTACGACCAGGGCTACACACGTGCTACAA<br>TGGCGCATACAAAGAGAAGCGACCTCGCGAGAGCAAGCGGACCTCATAA<br>AGTGCGTCGTAGTCCGGATTGGAGTCTGCAACTCGACTCCATGAAGTCGG<br>AATCGCTAGTAATCGTGGATCAGAATGCCACGGTGAATATGGTGTTCAATG<br>CTTTTCCCAGACGCGTGCTGAAGTCAAGCAAAGATGACGGGACCTACA |

**Supplementary Table 3. Strains used in this study.** The genotypes for each *E. coli* strain used are mentioned in the table.

| Species | Designation | Genotype / Strain |
|---|---|---|
| *Escherichia coli* | E. coli (MG1655) | MG1655 (K-12 F– λ– ilvG– rfb-50 rph-1) |
| | NEB Turbo | K-12 glnV44 thi-1 Δ(lac-proAB) galE15 galK16 R(zgb-210::Tn10)TetS endA1 fhuA2 Δ(mcrB-hsdSM)5(rK–mK–) F′[traD36 proAB+ lacIq lacZΔM15] |
| | MFDpir | MG1655 RP4-2-Tc::[ΔMu1::aac(3)IV-ΔaphA-Δnic35-ΔMu2::zeo] ΔdapA::(erm-pir) ΔrecA |

**Supplementary Table 4. Primers used for RT-qPCR and NGS.** The table includes the primer pairs used for RT-qPCR, along with the probe and reverse transcription primer where needed. The data for each primer pair are also noted for reference. The nucleotides in lower case refer to the Illumina adapters.

| Category | Target of the primer pair | Figure | Forward primer | | Reverse primer | |
|---|---|---|---|---|---|---|
| | | | Name | Sequence | Name | Sequence |
| RT-qPCR (dye) | Barcoded 16S rRNA | 2B, 3B, 3C, 4B, 4C | qRM17 | AGTCGGAATCGCTAGTAATCG | qRM14 | TGTAGGTCCCGTCATCTTTG |
| | 16S rRNA (native) | | qRM05 | CTAGCTGGTCTGAGAGGATG | qRM16 | TGTGCAATATTCCCCACTGC |
| RT-qPCR (probe) | Barcoded 16S rRNA | 5B | qRM17 | AGTCGGAATCGCTAGTAATCG | qRM14 | TGTAGGTCCCGTCATCTTTG |
| | | | qPCR probe | CGGTGAATATGGTGTTCAATGCTTTTCCC | | |
| Amplicon sequencing (NGS) | Barcoded 16S rRNA | 5C-F | 968F | AACGCGAAGAACCTTAC | LNK27A | GGTGTTCAATGCTTTTCCC |
| | 16S rRNA (native) | 5C-F | 968F | AACGCGAAGAACCTTAC | 1391R | GACGGGCGGTGWGTRCA |
| | Barcoded 16S rRNA (with adapter) | 5C-F | AOS20B | acactctttccctacacgacgctcttccgatctAACGCGAAGAACCTTAC | LNK28A | gactggagttcagacgtgtgctcttccgatctGGTGTTCAATGCTTTTCCC |
| | 16S rRNA (native) (with adapter) | 5C-F | | | DSZ05E | gactggagttcagacgtgtgctcttccgatctCGTTGACGGGCGGTGWGTRCA |
| | Reverse transcription primer | 5C-F | | | qRM14 | TGTAGGTCCCGTCATCTTTG |
| | | | | | 1391R | GACGGGCGGTGWGTRCA |

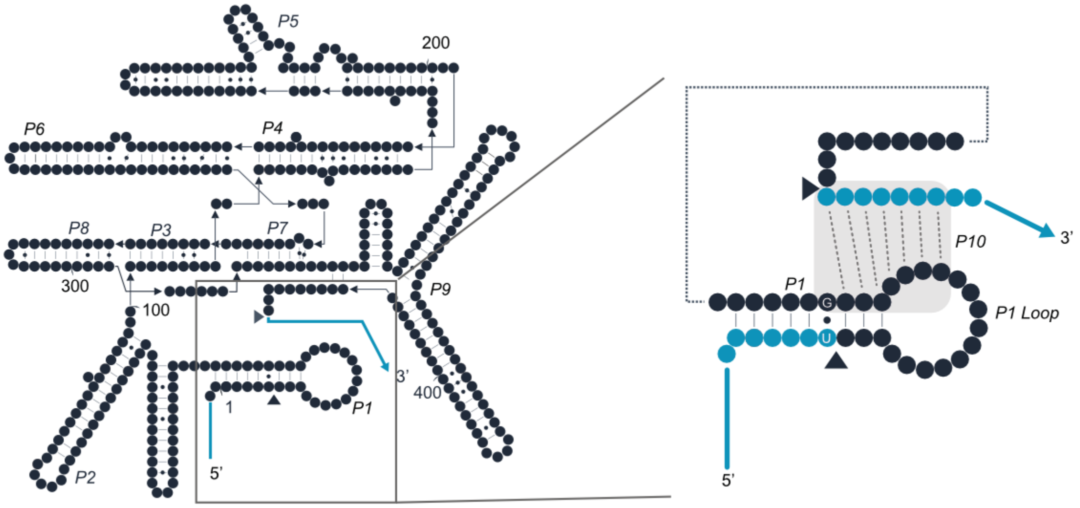


**Supplementary Figure 1. A schematic of the wild-type group I intron splicing ribozyme from *Tetrahymena thermophila*.** This ribozyme catalyses a cis-splicing reaction in which the intron (black) is removed from the flanking exons (blue; splice sites indicated by triangles) and joined into a single RNA strand. The 5' and 3' ends of the intron contain the P1 stem, the P1 loop, and the U•G wobble base pair necessary for splicing. The P10 interaction between the P1 and the 3' exon is also indicated in the right panel.

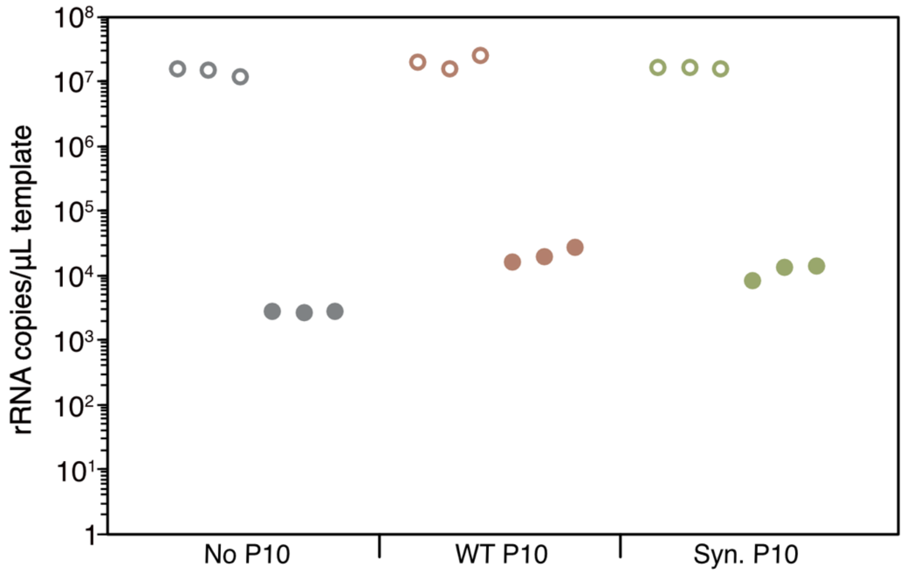


**Supplementary Figure 2**. **P10 interaction improves splicing efficiency**. Quantification of native and barcoded 16S rRNA using RT-qPCR in *E. coli* cells expressing RAM v1, WT P10, or Syn. P10. Empty data points represent three biological replicates for 16S rRNA, and solid data points represent three biological replicates for barcoded 16S rRNA.

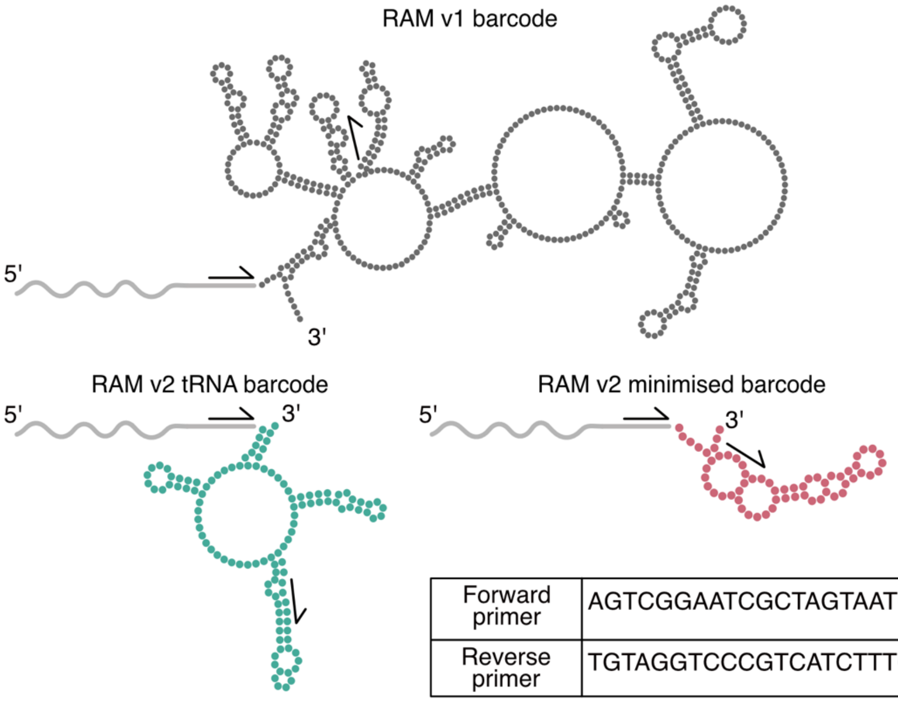


| Forward primer | AGTCGGAATCGCTAGTAATCG |
|---|---|
| Reverse primer | TGTAGGTCCCGTCATCTTTG |

**Supplementary Figure 3**. **Schematic of different RAM barcode designs**. Predicted secondary structure of different barcode designs (RAM v1, RAM v2 tRNA, and RAM v2 minimised) using the MXfold server, shown attached to 16S rRNA (grey). Primer binding sites used for RT-qPCR are indicated on the 16S rRNA and barcodes. Primer sequences are provided in the box.

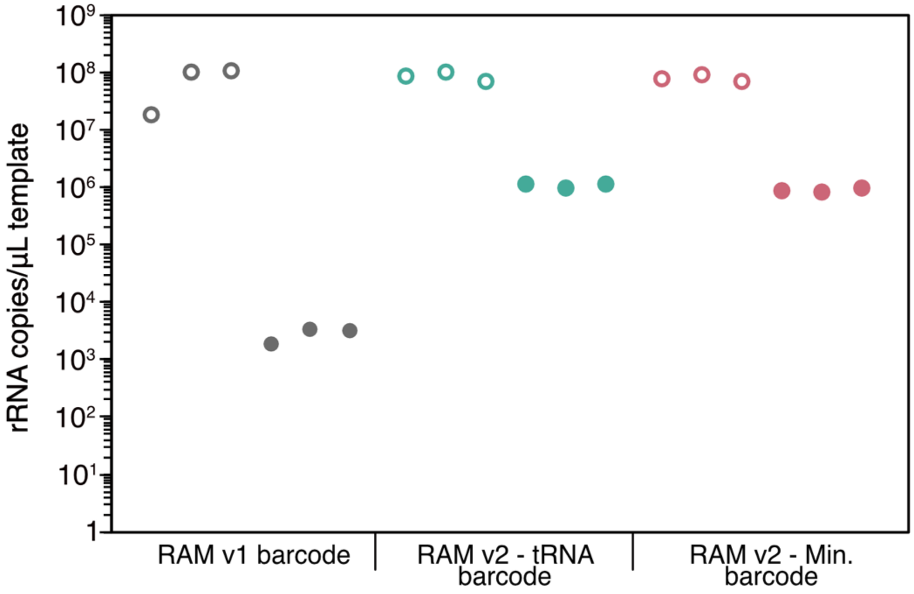


**Supplementary Figure 4**. **Barcode structure and length, combined with P10 interaction, increase the barcoded 16S rRNA signal.** Quantification of native and barcoded 16S rRNA using RT-qPCR in *E. coli* cells expressing RAM v1 or RAM v2 with the tRNA or minimised barcode. Empty data points represent three biological replicates for 16S rRNA, and solid data points represent three biological replicates for barcoded 16S rRNA.

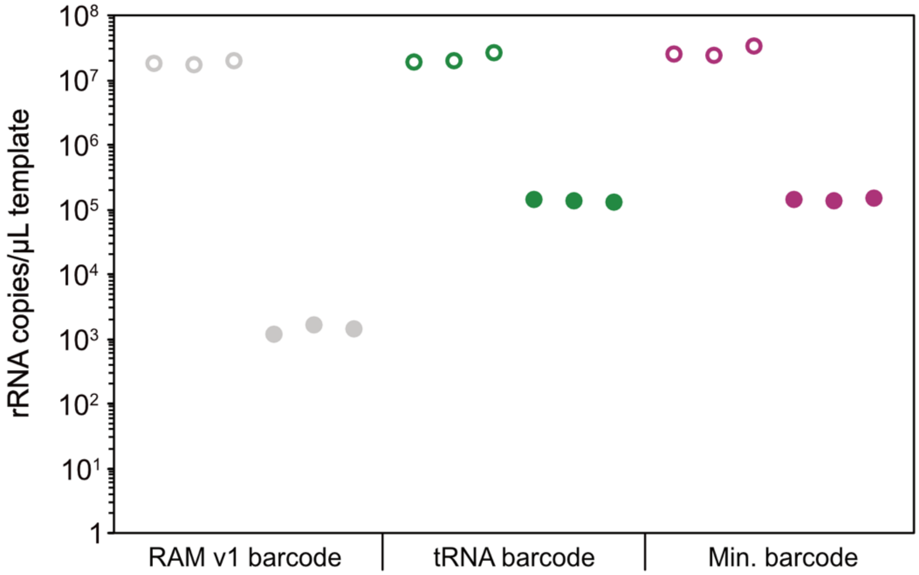


**Supplementary Figure 5**. **Barcode structure and length impact the signal of barcoded 16S rRNA.** Quantification of native and barcoded 16S rRNA in *E. coli* transformed with plasmids encoding RAM v1 or the two modified barcodes using RT-qPCR. Empty data points represent three biological replicates for 16S rRNA, and solid data points represent three biological replicates for barcoded 16S rRNA.

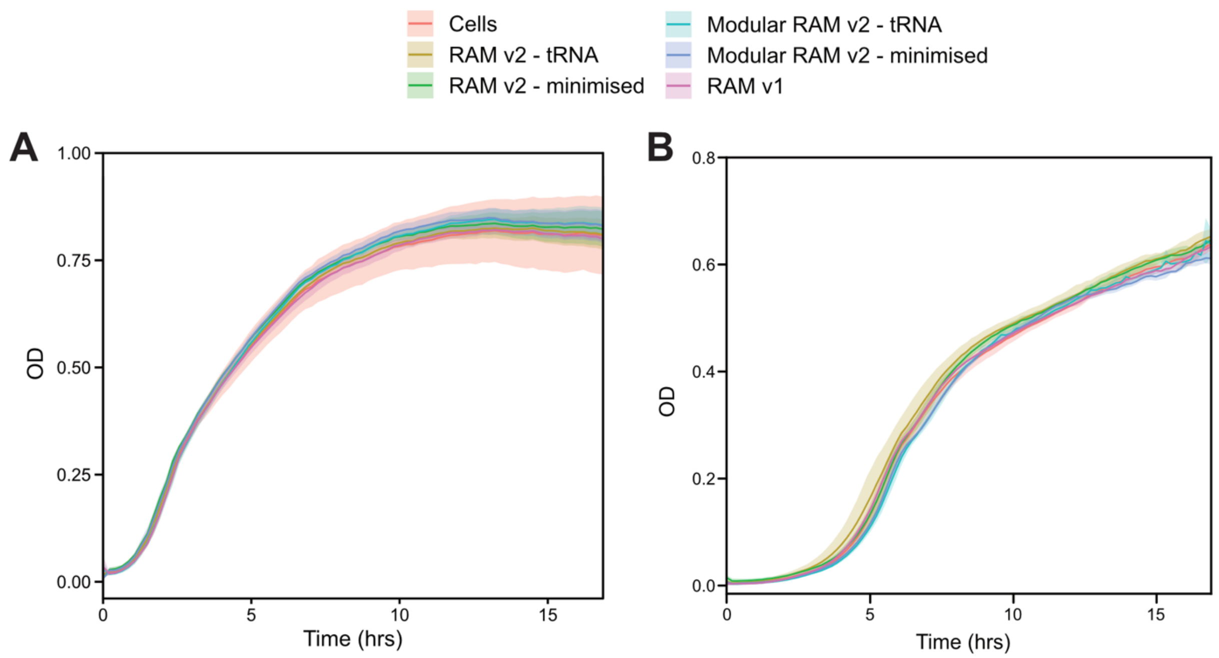


**Supplementary Figure 6**. **Rationally engineered RAM does not affect *E. coli* cell fitness.** Optical density (OD) measured at 600 nm of *E. coli* cells transformed with an empty control plasmid (Cells), the plasmid encoding RAM v1, RAM v2 with the two barcode designs, and modular RAM v2 with the two barcode designs, grown in (**A**) LB and (**B**) M9 media supplemented with casamino acids and thiamine. Solid lines represent the mean from 4 biological replicates with ±1 s.d. shown as shaded areas.

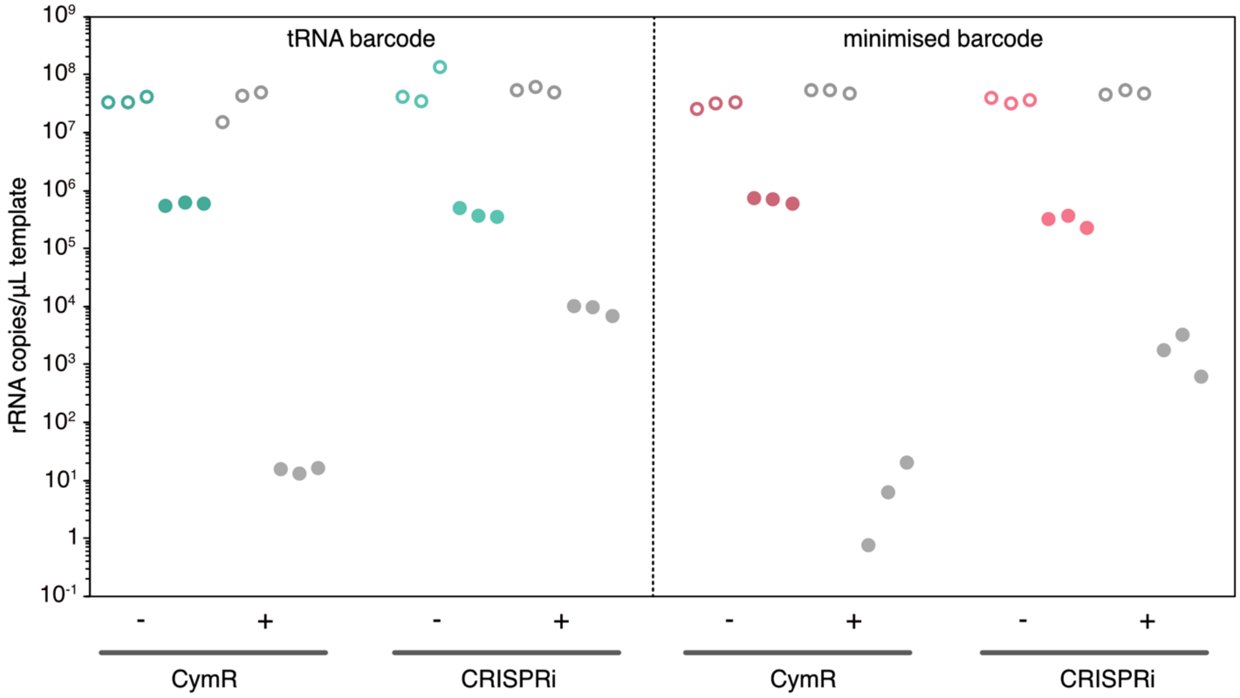


**Supplementary Figure 7**: **Repression of RAM and modular RAM in E.coli cells.** Quantification of native and barcoded 16S rRNA in *E. coli* containing plasmids encoding either non-modular or modular RAM v2, with and without the corresponding repression system present (CymR or CRISPRi) using RT-qPCR. Empty data points represent three biological replicates for 16S rRNA, and solid data points represent three biological replicates for barcoded 16S rRNA.

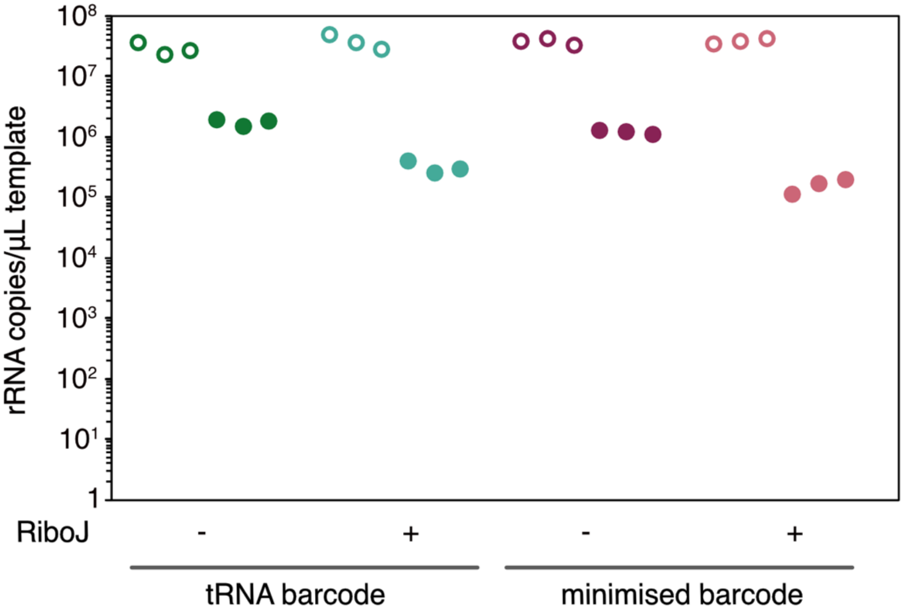


**Supplementary Figure 8**. **RiboJ in modular RAM v2 design results in a drop in barcoded 16S rRNA signal.** Quantification of native and barcoded 16S rRNA in RAM v2 constructs with and without the PlmJ insulator using RT-qPCR. Empty data points represent three biological replicates for 16S rRNA, and solid data points represent three biological replicates for barcoded 16S rRNA.

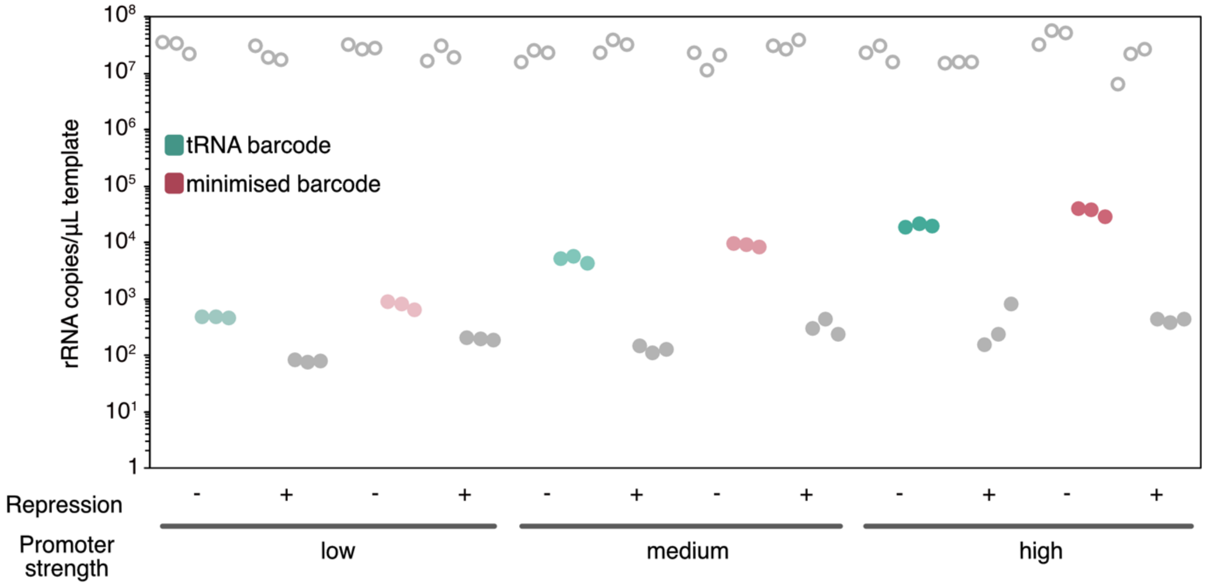


**Supplementary Figure 9. Modular RAM v2 works robustly across different promoter strengths.** Quantification of native and barcoded 16S rRNA in *E. coli* cells containing plasmids encoding the modular design using RT-qPCR. Three promoters of different strengths (J23105, J23101 and J23119) replaced the original promoter. Empty data points represent three biological replicates for 16S rRNA, and solid data points represent three biological replicates for barcoded 16S rRNA.

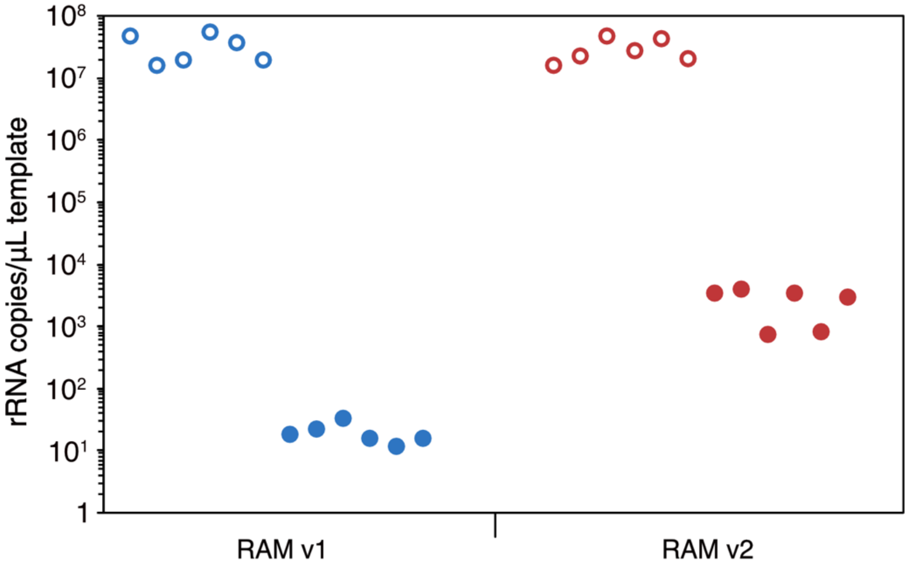


**Supplementary Figure 10**. **RAM v2 has increased barcode signal in a diverse Houston wastewater microbial community.** Quantification of native and barcoded 16S rRNA using RT–qPCR in a diverse Houston wastewater microbial community conjugated with plasmids encoding RAM v1 or RAM v2 - tRNA barcode. Empty data points represent six biological replicates for 16S rRNA, and solid data points represent six biological replicates for barcoded 16S rRNA.

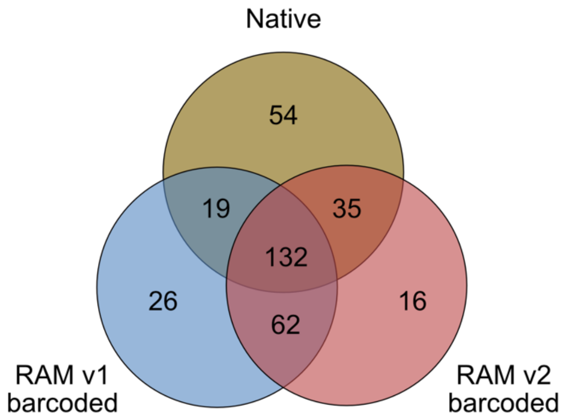


**Supplementary Figure 11. Native, RAM v1 barcoded, and RAM v2 barcoded ASVs observed in the wastewater microbial community.** The Venn diagram shows shared and unique ASVs in the native microbial community and transconjugant communities observed using RAM v1 and RAM v2. ASVs were determined via sequencing of 16S rRNA of 11 replicates for native ASVs (5 from RAM v1 and 6 from RAM v2 samples), and barcoded 16S rRNA of 5 replicates for RAM v1 and 6 replicates for RAM v2.

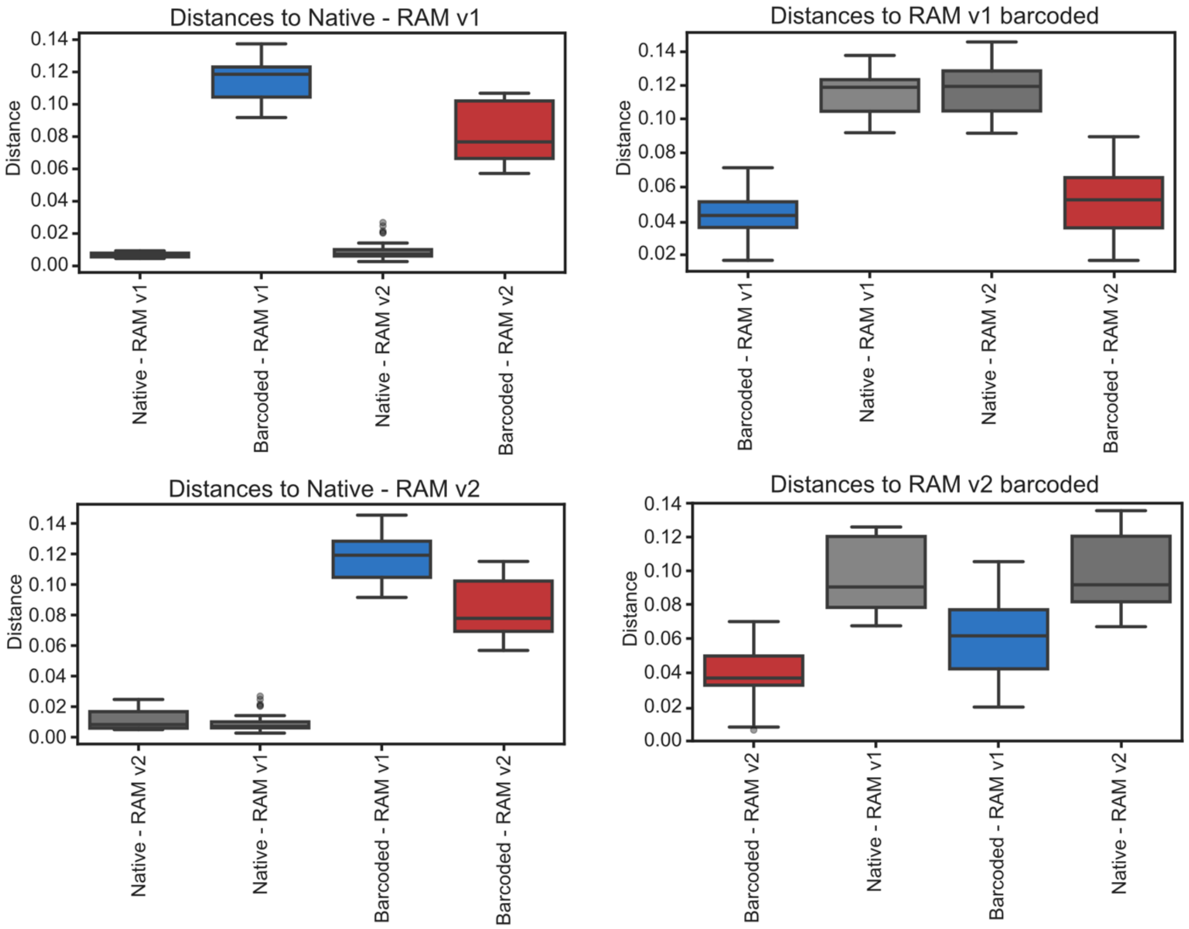


| Group 1 | Group 2 | Sample size | Permutations | pseudo-F | p-value | q-value |
|---|---|---|---|---|---|---|
| Native - RAM v1 | Barcoded - RAM v1 | 10 | 999 | 59.37 | 0.013 | 0.019 |
| Native - RAM v1 | Native - RAM v2 | 11 | 999 | 1.78 | 0.196 | 0.196 |
| Native - RAM v1 | Barcoded - RAM v2 | 11 | 999 | 45.56 | 0.003 | 0.008 |
| Barcoded - RAM v1 | Native - RAM v2 | 11 | 999 | 69.31 | 0.004 | 0.008 |
| Barcoded - RAM v1 | Barcoded - RAM v2 | 11 | 999 | 4.91 | 0.019 | 0.022 |
| Native - RAM v2 | Barcoded - RAM v2 | 12 | 999 | 52.37 | 0.002 | 0.008 |

**Supplementary Figure 12. PERMANOVA of a weighted unifrac distance matrix between the native and barcoded ASVs of RAM v1 and v2.** Four groups were analysed here: native ASVs in wastewater-RAM v1 conjugation, ASVs barcoded by RAM v1, native ASVs in wastewater-RAM v2 conjugation, and ASVs barcoded by RAM v2. The centre line in box plots represents the 50th percentile, the ends of the box represent the 25th and 75th percentiles of the box (interquartile range), and the lines represent the range of data points. The table shows statistics for each pairwise group comparison.

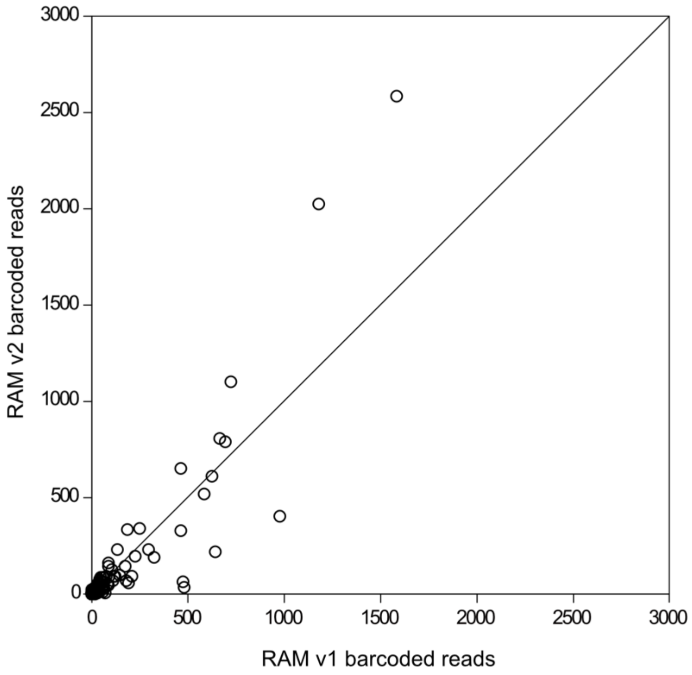


**Supplementary Figure 13. Scatter plot of ASVs barcoded by RAM v1 and RAM v2.** Read abundance for each ASV barcoded by both RAM v1 and v2. The solid line represents the 1:1 reference line, and the data points have an $R^2$ value of 0.8152.
